# WAVE complex self-organization templates lamellipodial formation

**DOI:** 10.1101/836585

**Authors:** Anne Pipathsouk, Rachel M. Brunetti, Jason P. Town, Artù Breuer, Patrina A. Pellett, Kyle Marchuk, Ngoc-Han T. Tran, Matthew F. Krummel, Dimitrios Stamou, Orion D. Weiner

## Abstract

How local interactions of actin regulators yield large-scale organization of cell shape and movement is not well understood. For example, why does the WAVE complex build lamellipodia, the broad sheet-like protrusions that power cell migration, whereas the homologous actin regulator N-WASP forms spiky finger-like actin networks? N-WASP is known to oligomerize into focal condensates that generate an actin finger. In contrast, the WAVE complex exhibits the linear distribution needed to generate an actin sheet. This linear organization of the WAVE complex could either arise from interactions with the actin cytoskeleton or could represent an ability of the complex to self-organize into a linear template. Using super-resolution microscopy, we find that the WAVE complex forms higher-order linear oligomers that curve into 270 nanometer-wide ring structures in the absence of actin polymer. These rings localize to the necks of membrane invaginations, which display saddle point geometries with positive curvature in one axis and negative curvature in the orthogonal axis. To investigate the molecular mechanism of saddle curvature enrichment, we show that the WAVE complex and IRSp53, a membrane curvature-sensitive protein, collaborate to recognize saddle curvature that IRSp53 cannot sense alone. This saddle preference for the WAVE complex could explain emergent cell behaviors, such as expanding and self-straightening lamellipodia as well as the ability of endothelial cells to recognize and seal transcellular holes. Our work highlights how partnering protein interactions enable complex shape sensing and how feedback between cell shape and actin regulators yields self-organized cell morphogenesis.

## INTRODUCTION

Cells manipulate the shape of their plasma membranes to execute a range of physiological functions from building the protrusions that drive cell motility to coordinating the membrane deformations that enable endocytosis. Actin polymerization plays a major role in coordinating these processes, but how cells specify the proper pattern of actin assembly to achieve these distinct shapes is unclear. Cells use nucleation-promoting factors (NPFs) to spatially and temporally control their patterns of actin polymerization (1,2). For example, neural Wiskott-Aldrich syndrome protein (N-WASP) and WASP-family verprolin homologous protein (WAVE) activate the actin-related protein 2/3 (Arp2/3) complex to seed actin nucleation (3,4). N-WASP and WAVE are nested in similar signaling topologies — both are stimulated by phosphoinositides, Rho GTPases, and curvature-sensitive Bin-amphiphysin-Rvs (BAR) domain proteins, both activate the Arp2/3 complex, both are recycled in an actin-dependent fashion, and both show evidence of oligomerization at the membrane (1,4–11) (**Fig 1A**). Despite these similarities, the structures they build are distinct: N-WASP participates in spiky filopodial protrusions, invadopodia, and endocytic vesicles (1,12–20), whereas WAVE participates in broad, sheet-like lamellipodial protrusions (21–24). The basis for this difference in NPF-induced cell morphology is not understood.

**Figure 1.**
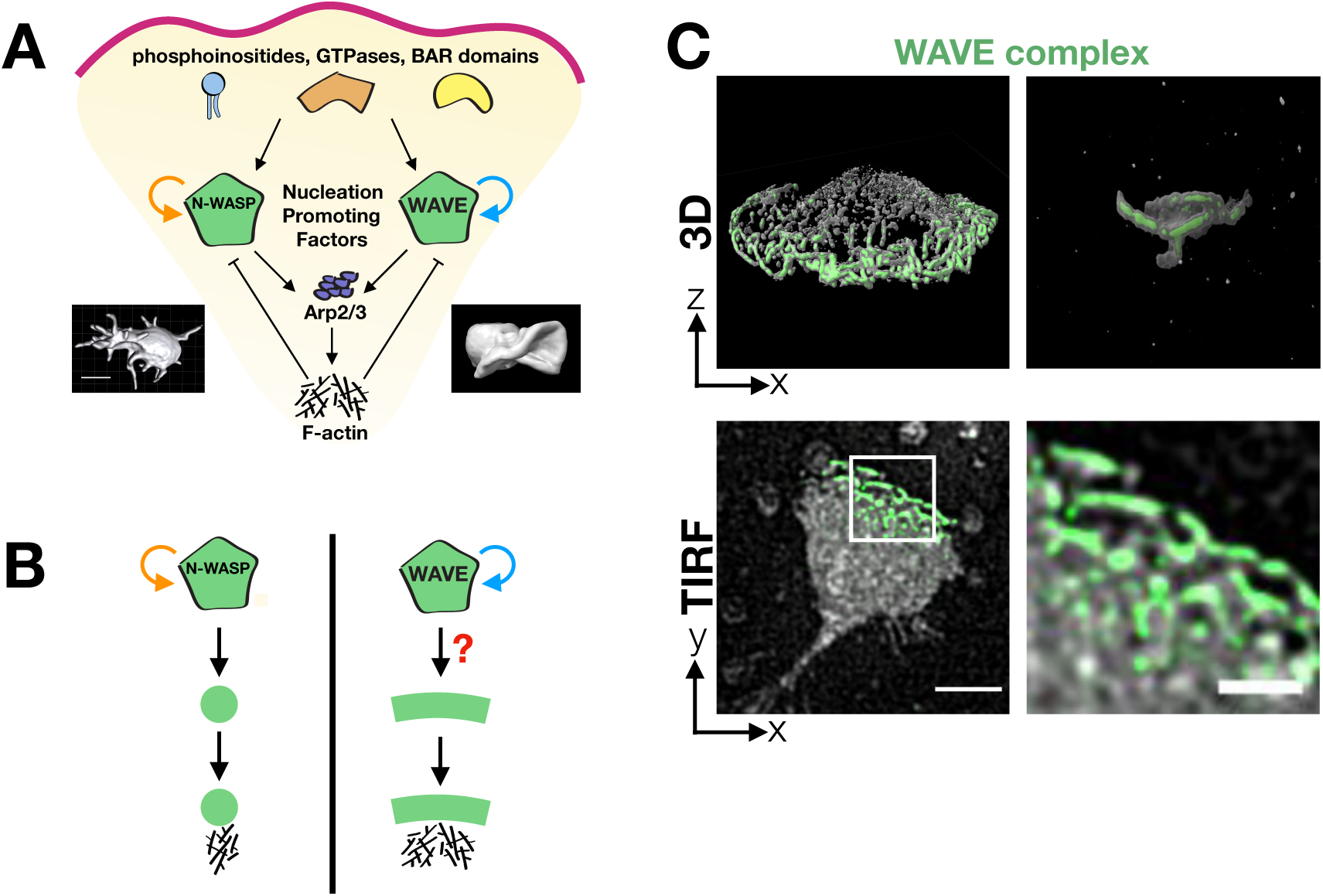
How does the WAVE complex achieve the linear spatial organization essential for lamellipodial generation? **(A)** Similar actin nucleation-promoting factors (NPFs) organize different cell morphologies. Given that the homologous NPFs WAVE complex and N-WASP are embedded in similar signaling cascades, how do they spatially organize different cellular morphologies? N-WASP canonically associates with filopodia, invadopodia, and endocytosis whereas the WAVE complex associates with lamellipodia. Images: left, chick cranial neural crest cell with multiple filopodia from (42) and right, head-on view of a neutrophil-like dHL60 lamellipodium (ChimeraX rendering of a confocal z-stack). **(B)** Schematic of how the NPF’s spatial organization could instruct the resulting actin morphologies. N-WASP’s positive feedback results in the focal organization expected for 1-dimensional finger-like actin structures. In order to build lamellipodia, does the WAVE complex’s positive feedback result in a linear organization to template 2-dimensional sheet-like actin structures? **(C)** Hem1-eGFP, a fluorescently tagged subunit of the WAVE complex, has a linear organization at tips of lamellipodia when dHL60 cells are stimulated with chemoattractant (10nM fMLP). Top: 3D imaging of the WAVE complex at tips of extending lamellipodia; left, widefield 3D reconstruction and right, lattice light sheet reconstruction of ruffles from a head-on view; see **S1 Video**. Bottom: WAVE complex’s linear organization viewed from the basal plasma membrane; simultaneous TIRF imaging of Hem1-eGFP (green) and membrane CellMask DeepRed dye (gray); see **S2 Video**. Scale bars: 5μm (left) and 2μm (right).

Biochemical reconstitutions have provided insights into how different patterns of actin nucleation lead to different morphological structures. Spatially unconstrained activation of the Arp2/3 complex produces filopodial-like structures on giant unilamellar vesicles (25). In contrast, spatially organizing Arp2/3 complex activation in a linear geometry (via glass rods (26) or UV micropatterned surfaces (27)) produces lamellipodial-like structures. These data suggest that while spiky, finger-like actin structures are the default morphology for the Arp2/3 complex activation on membranes, lamellipodium formation requires a linear structural template (28). What forms the basis of the linear template for lamellipodia in living cells is not known.

In cells, N-WASP forms the focal structure expected for finger-like protrusions (17,18,29), and the WAVE complex propagates as linear waves at the edges of lamellipodia (7,30). In the case of N-WASP, biochemical reconstitutions recapitulate its *in vivo* distribution: N-WASP and its multivalent binding partners generate biomolecular condensates (29,31). In the presence of the Arp2/3 complex and actin, these puncta-shaped condensates promote focal bursts of F-actin that produce spiky protrusions (31). In contrast, we do not know the basis of WAVE’s linear organization in cells (**Fig 1B**).

The WAVE complex is required for proper cell migration and shape maintenance across eukaryotes including mammals, amoeba, and plants (4,7,23,30,32–34). As a pentameric heterocomplex, the WAVE complex contains the subunits WAVE, Abi, HSPC300, Sra1/Cyfip1, and Nap1/Hem1 (22,35,36). In neutrophils and other motile cells, the dynamics of the WAVE complex closely corresponds to the leading edge’s morphology and pattern of advance (7). The WAVE complex is required for the formation of lamellipodia and efficient chemotaxis (directed cell migration); previous work found that siRNA knockdown and CRISPR-Cas9 gene inactivation of the WAVE complex in neutrophil-like differentiated HL60 cells (dHL60s) and other immune cells results in defective chemotaxis and lamellipodia formation (22–24,37–39).

Here, we investigate how the WAVE complex achieves the linear distribution essential for lamellipodial actin organization. We found that, in the absence of actin polymer, the WAVE complex forms nanoscale oligomeric ring structures that associate with regions of membrane that have saddle point curvature. To further explore saddle curvature enrichment, we analyzed the WAVE complex’s localization to transendothelial cell macroaperture (TEMs) tunnels, which are transcellular holes that leukocytes and various pathogens generate in endothelial cells to cross tissue barriers (40). As the endothelial cell seals the hole, the TEM displays persistent saddle curvature (41). We found that the WAVE complex enriches to closing TEMs, which suggests it may have a role in linking the recognition to the sealing of TEMs. To identify a potential mechanism of membrane curvature recognition, we explored the WAVE complex’s partnering interactions with IRSp53, an inverse BAR domain protein, and found that their collaboration is required for IRSp53 to recognize lamellipodial and saddle curvature geometries. We propose that the WAVE complex’s nanoscale, oligomeric assembly could explain emergent, self-organizing behaviors of cell morphogenesis including expanding and self-straightening lamellipodia and the recognition and closure of transcellular holes.

## RESULTS

In neutrophil-like dHL60 cells and a range of other motile cells, the WAVE complex can be seen propagating as a linear “wave” structure at the edges of membrane ruffles and lamellipodia (**Fig 1C**). The WAVE complex’s propagating dynamics are thought to account for many behaviors of motile cells, such as continuous extending fronts and the ability to migrate around barriers (7). This dynamic propagation arises from an excitable feedback network with positive feedback (WAVE recruits more WAVE) and negative feedback loops (WAVE stimulates actin polymerization, which inhibits WAVE association with the membrane) (7,43). One source of negative feedback appears to be the force of actin polymerization that strips WAVE off the plasma membrane (43). In contrast, the mechanism and spatial organization of positive feedback, i.e. WAVE self-association, are not well understood. We were particularly interested in understanding whether the linear patterns of the WAVE complex in lamellipodia are dependent on an interaction between these positive and negative feedback loops, as is the case for other excitable networks (44), or whether the WAVE complex’s self-association has a specific geometric organization (**Fig 2A**). We can use inhibitors of actin polymerization to deplete F-actin to distinguish the two models and assay whether the WAVE complex maintains a stereotyped linear structure independently of F-actin.

**Figure 2.**
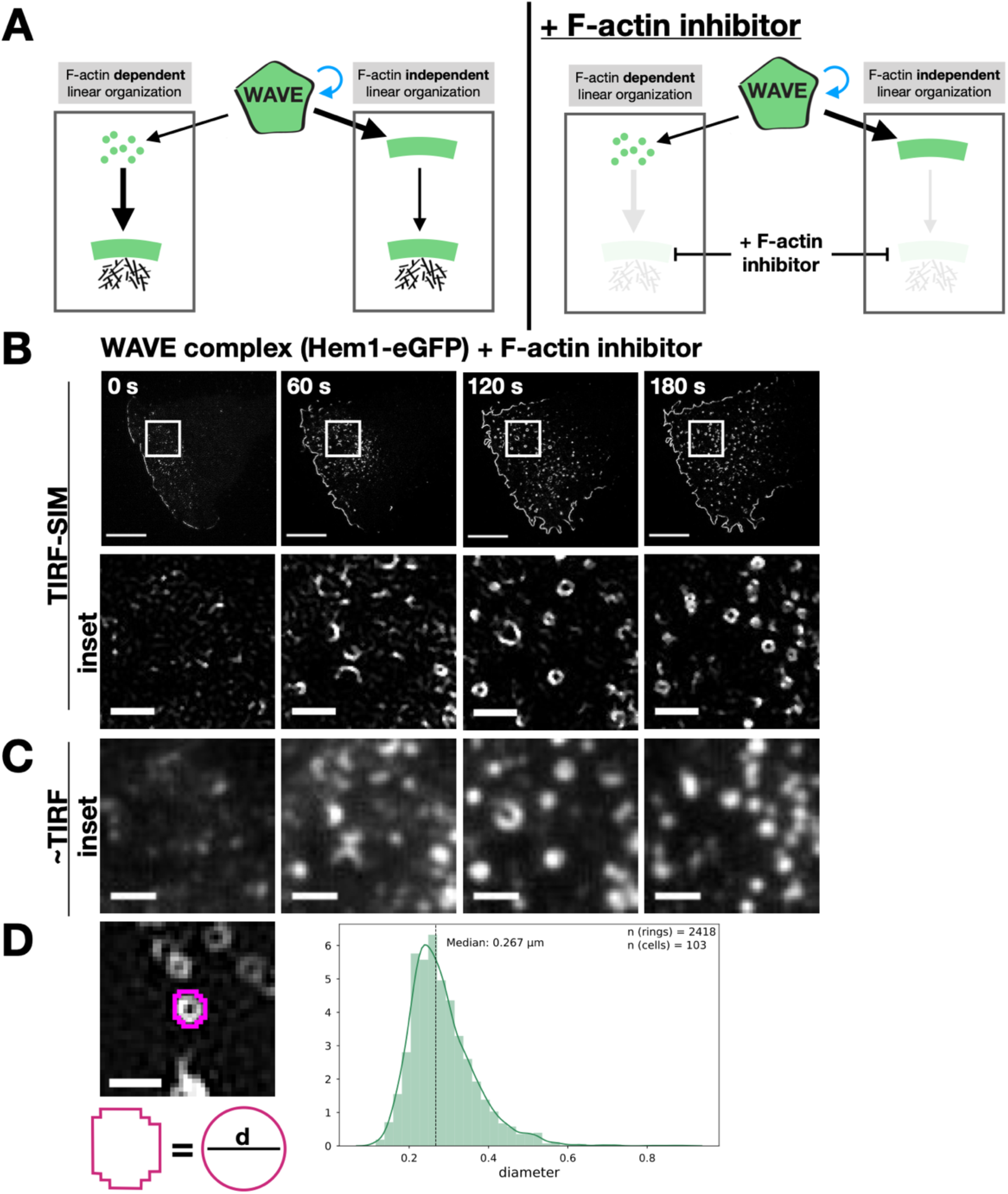
WAVE complex organizes as nanoscale rings in the absence of the actin cytoskeleton. **(A)** Models of the basis of the WAVE complex linear organization and the experimental design to distinguish the models. Left: the WAVE complex could achieve its linear organization at the lamellipodial edge in a manner that is dependent on its interactions with the actin cytoskeleton (left) or that is established independent of the actin cytoskeleton (right). Right: addition of a Factin inhibitor can be used to distinguish these models. **(B)** Super-resolution total internal reflection fluorescence structured illumination microscopy (TIRF-SIM) imaging reveals that the WAVE complex forms ring structures in the absence of actin polymers. dHL60 cells expressing Hem1-eGFP were acutely stimulated with latrunculin A (5μM final); see **S3 Video**. Bottom row are the white insets. Time in seconds; scale bars: 5μm (top) and 1μm (insets). **(C)** Conventional TIRF resolution comparison highlights the need for super-resolution microscopy to resolve diffraction-limited WAVE complex puncta as rings. Conventional resolution TIRF images were created by sum-projecting the 9 images (3 phases * 3 angles) that construct TIRF-SIM images of (**B**); see **S4 Video.** Scale bar: 1μm. **(D)** The median diameter of the WAVE complex ring structures is 267 nanometers. Left, example measurement of fitting the ring’s perimeter to a perfect circle and calculating the circle’s diameter, d. Right, histogram of diameters of rings across a range of F-actin inhibiting drugs and concentrations of n = 2418 rings from 103 cells; see **S2 Fig**.

When we previously visualized the pattern of the WAVE complex organization (via fluorescently-tagged hematopoietic protein 1 [Hem1], a subunit of the WAVE complex) in the absence of the actin cytoskeleton, standard total internal reflection fluorescence (TIRF) microscopy showed amorphous punctate structures of the WAVE complex (7). For our current study, we revisited this experiment with super-resolution microscopy, which enables us to investigate any previously diffraction-limited organization of the WAVE complex. We leveraged TIRF-structured illumination microscopy (TIRF-SIM), a super-resolution technique that enables a 2-fold increase in resolution in the TIRF plane. When the F-actin inhibitor latrunculin A was acutely added to dHL60 cells to deplete F-actin, TIRF-SIM imaging of Hem1-eGFP revealed that the WAVE complex organizes into highly stereotyped 270nm-wide ring structures (**Fig 2B-D**). Importantly, super-resolution microscopy was required to resolve the nanometer-scale ring structures because they otherwise appear as amorphous blob-like structures by conventional TIRF (**Fig 2C**).

We observed two modes of ring generation: either *de novo* formation by recruitment of new WAVE complex to the plasma membrane following actin depolymerization or redistribution of membrane-bound WAVE complex from lamellipodia that locally “collapsed” into rings (**S1 Fig**). The WAVE complex rings were observed across a range of F-actin inhibitors and concentrations (**S2 Fig**), super-resolution modalities (**S3A Fig**), tagged fluorescent proteins (**S3B Fig**), cell types, and specific WAVE complex subunits (**S3C Fig**). Since latrunculin treatment results in lowered plasma membrane tension (45), the rings may represent the favored oligomeric organization of the WAVE complex when freed from the constraints of the cytoskeleton and tension in the plasma membrane. These experiments suggest that the WAVE complex’s positive feedback forms nanoscale, oligomeric rings.

Next, we wondered how these rings are organized relative to the morphology of the membrane. During cell migration, the WAVE complex is closely associated with the propagating edge of lamellipodial protrusions (**Fig 1C**). When imaging the WAVE complex and the plasma membrane in the absence of F-actin, we found that the WAVE complex rings localized to the boundary where coverslip-opposed membrane leaves the TIRF field (**Fig 3A**). This membrane distribution suggests an association to the necks of membrane invaginations (**Fig 3B**), and electron microscopy experiments are consistent with this membrane geometry (**S4 Fig**).

**Figure 3.**
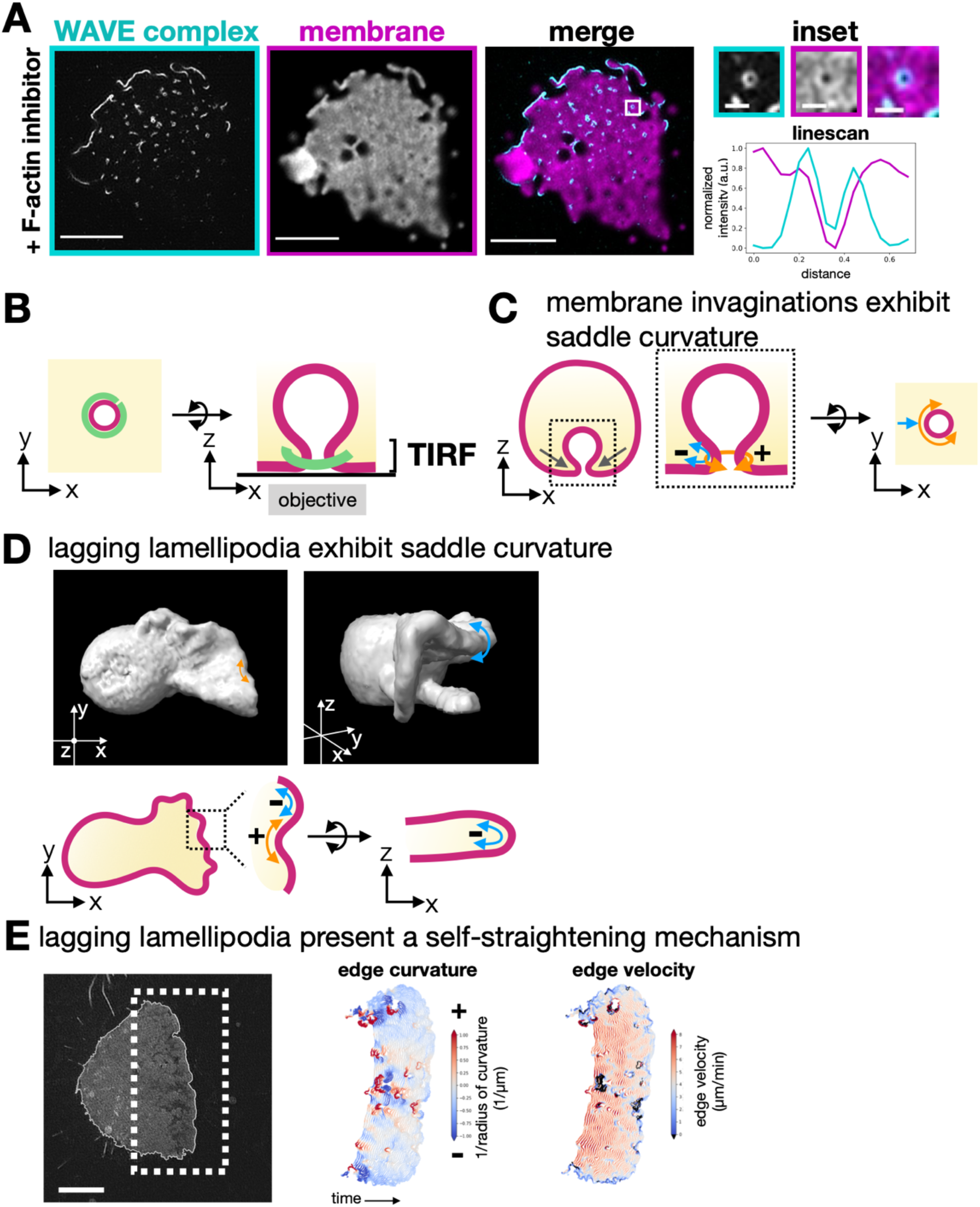
WAVE complex rings associate with membrane saddle points. **(A)** TIRF-SIM imaging of the WAVE complex (Hem1-eGFP; cyan) reveals rings localized at the boundary of where the plasma membrane (DiD-labeled membrane; magenta) exits the TIRF plane; see **S5 Video**. This membrane organization suggests enrichment to the necks of membrane invaginations. Right, graph of a linescan across a ring. Scale bars: 5μm and 500nm (inset). **(B)** Schematic of proposed organization of the WAVE complex ring structure (green) around the neck of a membrane invagination (magenta) as seen from the top, XY plane (left), or the side, XZ plane (right). **(C)** Schematic of a membrane invagination’s saddle curvature. The necks of invaginations display saddle geometry of positive (orange; the curve around the invagination neck) and negative (blue; the curve perpendicular to the invagination neck) curvatures. **(D)** Lagging portions of lamellipodia display saddle geometry. Top: ChimeraX rendering of a confocal z-stack membrane-bound marker in dHL60s. Top left: view of the top of the cell shows lagging portions of a lamellipodium extension. Top right: tilted orthogonal view highlights the negative curvature and sheet-like morphology of the lamellipodium. Bottom: schematic of curvatures of a lamellipodium. **(E)** Lagging lamellipodia present a self-straightening mechanism. Left: TIRF-SIM image of membrane signal with its membrane edge outlined. Scale bar: 5μm. Middle: curvature analysis of the advancing front edge over time (1 frame every 2 seconds). Cool-warm scale indicates negative-positive curvature. Right: velocity analysis of the advancing front edge over time. Areas of positive curvature that become less positive, i.e. self-straighten, are associated with areas of increasing velocity.

WAVE complex’s propensity to enrich around the necks of membrane invaginations may give insight into the WAVE complex’s membrane geometry preferences. The geometry of the necks of membrane invaginations from the cytosolic/WAVE complex’s perspective consists of saddle points, defined by principal curvatures that are positive in one axis (the curve around the invagination neck) and negative in the other axis (the curve perpendicular to the invagination neck) (**Fig 3C**). Saddle geometries are not only present at membrane invaginations but also at areas of the protruding lamellipodium that transiently fall behind or “lag” (**Fig 3D**). As the membrane extends, it is laterally non-uniform and displays positive and negative curvatures (**Fig 3D** left) while it maintains an axial, sheet-like negative curvature (**Fig 3D** right). In order to maintain a coherent protruding lamellipodium, the lagging portion, which exhibits positive curvature, must become less positively curved, i.e. straighten, and accelerate to “catch up” (**Fig 3E**). This acceleration could be mediated by a localized increase in F-actin polymerization, via WAVE complex NPF activity, to push and straighten the lagging regions of the lamellipodium. The result of these interactions would be a homeostatic self-straightening mechanism for a saddle-enriching WAVE complex.

To further explore the role that saddle curvature recognition may have in the emergent control of cell shape, we took advantage of transendothelial cell macroaperture (TEM) tunnel physiology. As leukocytes undergo diapedesis out of the blood vessel, they can migrate either in between endothelial cells or they can generate a TEM, a transcellular hole, to migrate through an endothelial cell (40). The hole is formed from fusing the apical and basal plasma membrane of the endothelial cell (46). To heal the transcellular hole and prevent pathogen dissemination, the affected endothelial cell has mechanisms to seal the TEM (47–49). The sealing process requires the recognition of TEMs, which exhibit saddle curvature, and subsequent closure of the hole (**Fig 4A-B**) (47). TEM closure is mediated by actin and the Arp2/3 complex dependent mechanisms (**S5 Fig**), but the relevant Arp2/3 complex activator has not been identified (47,50). Closure has been characterized as lamellipodial-like (48), where the height of the TEM edge is ∼100nm by atomic force microscopy measurements (47). As TEMs close, the negative curvature of the lamellipodia-like extension remains constant while the positive curvature around the TEM increases (fractions of microns^-1^ to microns^-1^) (47,51). To generate TEMs, we inhibited actomyosin contractility, which is also the mechanism that some pathogens use to generate TEMs (40,47,48,50). For this purpose, we studied HUVECs (human umbilical vein endothelial cells) treated with ROCK inhibitor Y27632. In this experimental setup, we imaged the WAVE complex (via fluorescently-tagged Nap1, a homologue of Hem1) and found that it also enriched to TEMs (**Fig 4C**). In addition to localizing to the necks of membrane invaginations in F-actin inhibited dHL60 cells, the WAVE complex’s TEM association represents another example of saddle enrichment (**Fig 4D**).

**Figure 4.**
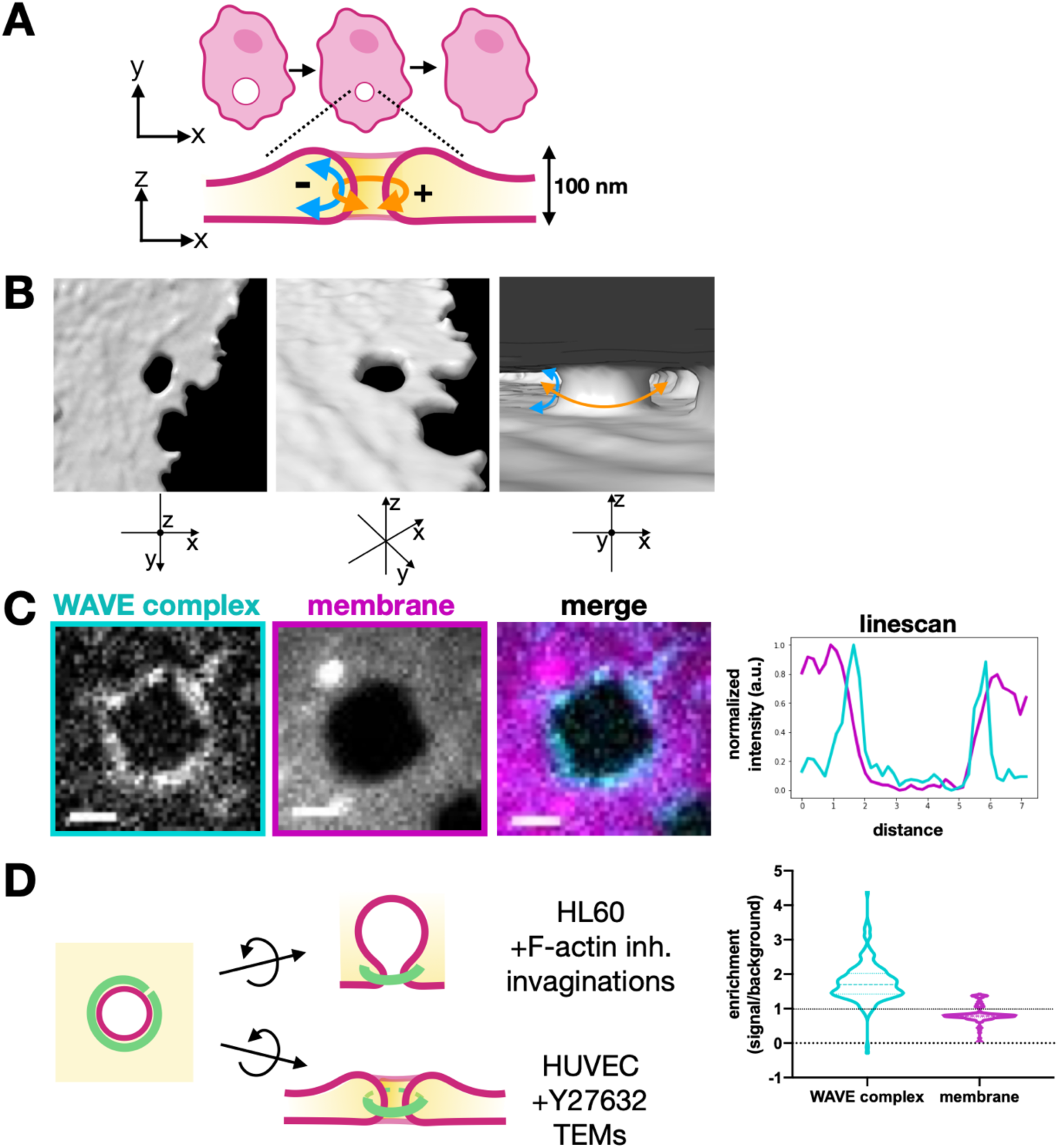
WAVE complex enriches to transendothelial cell macroaperture (TEM) tunnel saddle points. **(A)** Schematic of TEM closure and its saddle geometry. **(B)** ChimeraX rendering of spinning disk confocal imaging of a fixed HUVEC cell treated with Y27632 (50μM) and labeled with a membrane dye (CellMask DeepRed). Middle: tilted view. Right: view from inside the cell facing a TEM and its saddle geometry. See **S6 Video**. **(C)** HUVEC cells expressing eGFP-tagged Nap1, a WAVE complex subunit, and stained with membrane dye CellMask DeepRed show the WAVE complex localized at a TEM. Spinning disk confocal imaging, scale bar: 2μm. Top graph: linescan across the TEM; WAVE complex (cyan) and membrane dye (magenta). Bottom graph: violin plot of enrichment, which was measured as the ratio between the signal intensity per unit area at the TEMs compared to the background intensity per unit area. A value of 1 indicates no enrichment at the TEMs (dotted line), and a value above 1 indicates enrichment. Each time point throughout TEM closure was considered a single data point; WAVE complex and membrane both had n = 178 (from 6 TEMs) from at least 3 independent experiments per condition. **(D)** Schematic showing the WAVE complex localization to saddle geometries of membrane invaginations in F-actin-inhibited dHL60s as well as TEMs in Y27632-treated HUVECs.

Next, we sought to understand how the WAVE complex recognizes saddle geometries. As saddles have both negative and positive curvatures, there may be two domains that sense the different curvatures. Though the WAVE complex has membrane-binding motifs (6), it lacks any well-characterized curvature sensing motifs. However, the WAVE complex does directly interact with IRSp53 (insulin receptor tyrosine kinase substrate protein of 53 kDa; human ortholog is BAIAP2 [brain-specific angiogenesis inhibitor 1-associated protein 2]), a member of the inverse BAR (I-BAR) domain family of curvature-sensitive proteins (10,11,52). I-BAR domains dimerize into a convex geometry that binds phosphoinositide-rich membranes and are the canonical negative curvature sensing motif that associates with lamellipodia and filopodia (53–56). IRSp53 has an I-BAR domain at its N-terminus and a number of protein-protein interaction domains including a Src homology 3 (SH3) domain that binds to the WAVE complex’s proline-rich domain (10,52,57) (**Fig 5A**).

**Figure 5.**
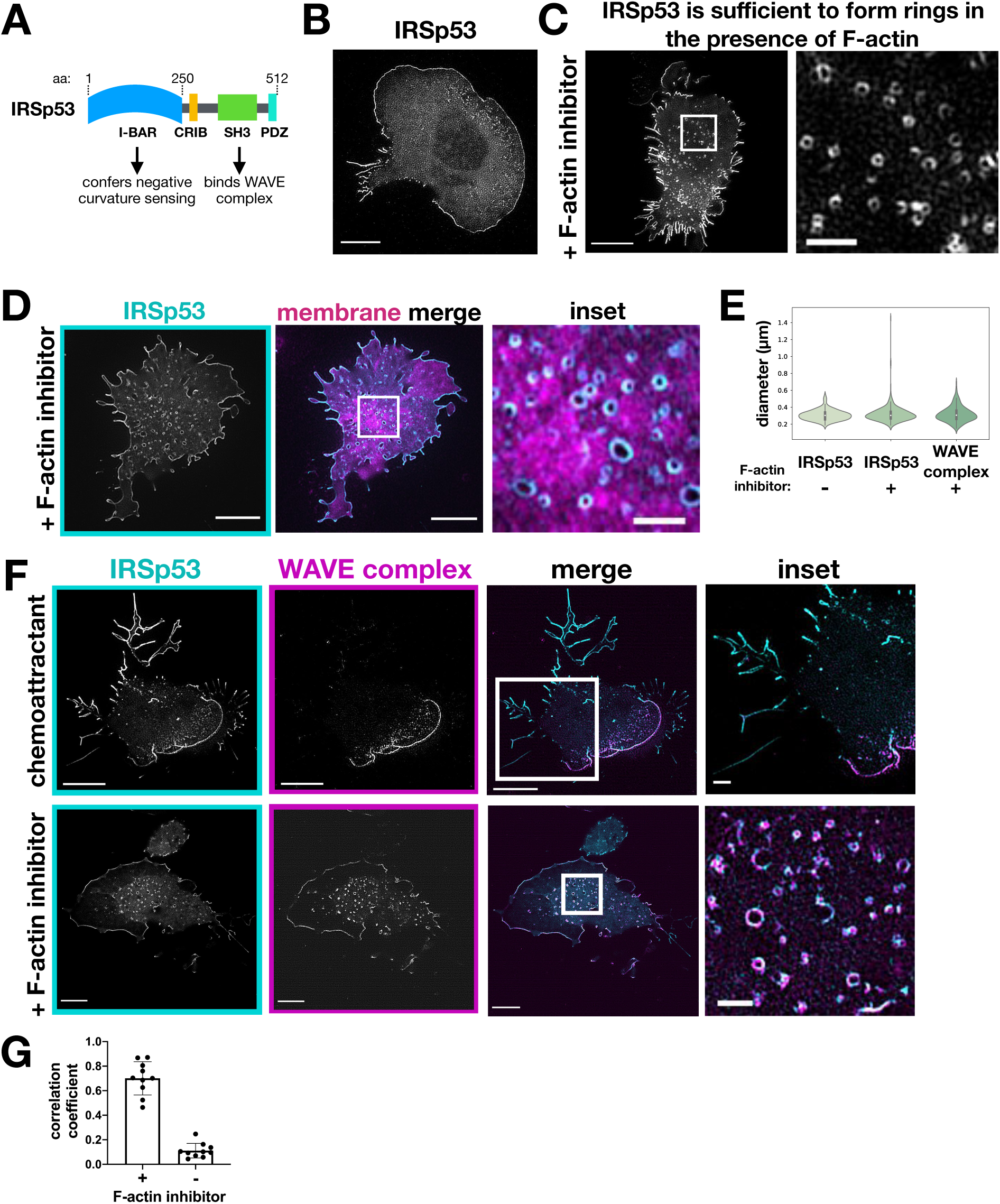
IRSp53 and the WAVE complex colocalize at lamellipodia and regions of saddle curvature but not at filopodia-like structures. **(A)** Domain structure of IRSp53. The I-BAR domain confers negative membrane curvature sensing and the SH3 domain enables binding to the WAVE complex. **(B)** dHL60 cell expressing IRSp53-eGFP and stimulated with chemoattractant (10nM fMLP) show IRSp53 localization to both lamellipodia and filopodia-like structures. TIRF-SIM imaging, scale bar: 5μm. **(C)** dHL60 cell with high expression of IRSp53-eGFP stimulated with chemoattractant (10nM fMLP) shows that IRSp53 is sufficient to form ring structures even in the presence of an intact actin cytoskeleton; see **S8 Video.** TIRF-SIM imaging, scale bars: 5μm (left) and 1μm (inset). **(D)** dHL60 cell treated with F-actin inhibitor (500nM latrunculin B) shows IRSp53-eGFP (cyan) forms ring structures. Right, membrane merge (CellMask DeepRed; magenta). TIRF-SIM imaging, scale bars: 5μm (left) and 1μm (inset). **(E)** Graph comparing the diameters of the WAVE complex rings with F-actin inhibition and IRSp53 rings with (+) or without (-) F-actin inhibition (500nM latrunculin B). Diameters measured in the same fashion shown in **Fig 2D**. Violin plot of rings; IRSp53 without F-actin inhibition n = 157 from 10 cells, IRSp53 with F-actin inhibition n = 483 from 10 cells, the WAVE complex n = 202 from 13 cells; cells pooled from at least 3 independent experiments per condition; Kruskal-Wallis test, nonsignificant P value > 0.05. **(F)** IRSp53 and the WAVE complex colocalize at lamellipodia (top) and ring structures following actin depolymerization (bottom). IRSp53 also localizes to filopodia-like structures whereas the WAVE complex is excluded from those regions (top). Cells expressing both IRSp53-eGFP and Hem1-mCherry were treated with chemoattractant (10nM fMLP, top; see **S9 Video**) or F-actin inhibitor (500nM latrunculin B, bottom). TIRF-SIM imaging; scale bars: 5μm and 1μm (inset). **(G)** Graph measuring colocalization between IRSp53-eGFP and Hem1-mCherry in conditions with or without F-actin inhibition. Barplot shows mean +/-standard deviation of Manders M1 correlation coefficient (fraction of IRSp53 in compartments containing Hem1); (+) F-actin inhibition, mean = 0.70 +/-0.14, n = 10 cells, (-) F-actin inhibition, mean = 0.11 +/-0.06, n = 10 cells; cells pooled from at least 3 independent experiments of TIRF-SIM imaging.

To explore the range of geometries sensed by IRSp53, we imaged IRSp53-eGFP in dHL60 cells. Full-length IRSp53 localized to multiple areas of negative curvature: lamellipodia (10,11,52,58,59), filopodia-like tubules (58–61), and rings (**Fig 5B-C**). Importantly, high expression of IRSp53 is sufficient to form ring structures even without F-actin inhibition. When cells were treated with a F-actin inhibitor, IRSp53 also localized to ring structures (**Fig 5D**). IRSp53 rings formed with or without F-actin present had comparable diameters to the WAVE complex rings formed without F-actin present (**Fig 5E**). This characteristic size suggests a preferred geometry for IRSp53 and the WAVE complex’s membrane recruitment in cells.

To probe whether the WAVE complex follows IRSp53 to sites of negative curvature, we imaged IRSp53 and the WAVE complex in the same cell (via IRSp53-eGFP and Hem1-mCherry). IRSp53 and the WAVE complex only co-localized at protruding lamellipodia during cell migration (**Fig 5F-G**). In addition to lamellipodial enrichment, IRSp53 also localized to filopodia-like tubules, whereas the WAVE complex was restricted to the lamellipodium. These data suggest that the WAVE complex does not follow IRSp53 to all sites of negative curvature. When cells were treated with a F-actin inhibitor, both the WAVE complex and IRSp53 co-localized to ring structures (**Fig 5F-G**), consistent with both proteins showing saddle curvature enrichment. While IRSp53 localized to multiple sites displaying negative curvature, the WAVE complex was specifically enriched to lamellipodia and regions of saddle curvature.

Since the WAVE complex and IRSp53 colocalize in a subset of cell structures, we wondered whether IRSp53’s localization pattern is partially dependent on its interactions with the WAVE complex. IRSp53 consists of several functional domains that could contribute to its localization, in particular an I-BAR domain that senses negative curvature and an SH3 domain that interacts with the WAVE complex. To investigate how these domains contribute to the overall pattern of IRSp53 enrichment, we generated two structure-function constructs tagged with eGFP at the C-terminus: an “I-BAR” only domain that consisted of IRSp53’s first 250 amino acids and a “C-term” only construct that lacks the I-BAR domain but contains the region that interacts with the WAVE complex (**Fig 6A**). While full-length IRSp53 enriched to both lamellipodia and filopodia-like structures, the I-BAR domain preferentially enriched to filopodia-like structures, and the C-term domain remained cytosolic (**Fig 6B**). In latrunculin-treated cells, full-length IRSp53 formed rings while the I-BAR and C-term constructs failed to enrich as ring structures (**Fig 6C**). These data suggest that IRSp53 requires both its I-BAR domain and its C-terminal portion, containing its SH3-WAVE complex interacting domain, to properly enrich to lamellipodia and saddle points.

**Figure 6.**
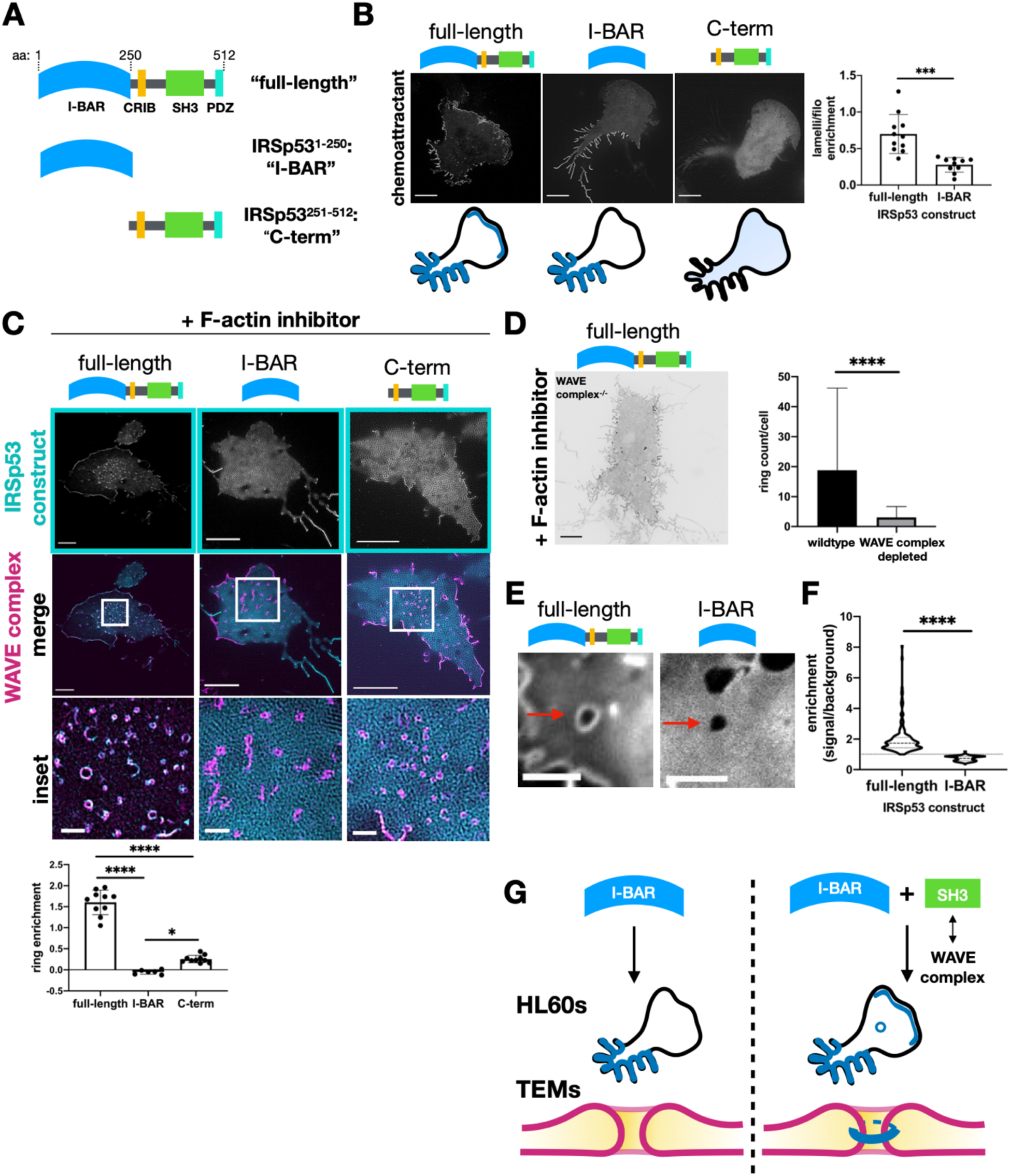
IRSp53 requires both its I-BAR domain and its interactions with the WAVE complex to localize to lamellipodia and regions of saddle curvature. **(A)** Schematic of IRSp53 structure-function constructs. **(B)** Chemoattractant stimulated dHL60s expressing the eGFP-tagged IRSp53 constructs. Full-length IRSp53 enriches to lamellipodia and filopodia-like structures; I-BAR enriches to filopodia-like structures but not to lamellipodia; and C-term is cytosolic. Summary of the IRSp53 construct localization below. TIRF-SIM imaging; scale bar: 5μm. Right, graph of IRSp53 enrichment ratio of signal at lamellipodia per unit area vs signal at filopodia per unit area for full-length and I-BAR IRSp53 constructs. Graph shows mean +/-standard deviation; full-length mean = .70 +/-0.27, n = 11 cells; I-BAR mean = 0.28 +/-0.10, n = 10 cells; cells pooled from at least 3 independent experiments per condition; unpaired t-test two-tailed ***P value <0.001. **(C)** F-actin inhibited dHL60s expressing IRSp53 and the WAVE complex. Middle row shows IRSp53-eGFP construct (cyan) overlay with Hem1-mCherry (magenta) rings. The I-BAR and C-term constructs fail to enrich robustly as ring structures. TIRF-SIM imaging; scale bars: 5μm and 1μm (inset). Graph comparing the signal enrichment of IRSp53 constructs’ signal per unit area of rings (defined by Hem1-mCherry) over the background per unit area. Graph displays the mean +/-standard deviation where all rings within a cell were aggregated; full-length mean = 1.6 +/-0.29, n = 10 cells from 3 independent experiments, I-BAR mean = -.05 +/-0.05, n = 6 cells from 2 independent experiments, C-term mean = 0.25 +/-0.09, n = 10 cells from 2 independent experiments; one-way ANOVA with two-tailed P value <0.0001 with Tukey’s multiple comparisons follow-up tests, P values **** <0.0001, * <0.05. **(D)** Full-length IRSp53 fails to enrich as ring structures in the absence of the WAVE complex. Inverted display of IRSp53-eGFP expressed in a WAVE complex depleted cell treated with latrunculin B (500nM). TIRF-SIM imaging; scale bar: 5μm. Graph comparing the mean +/-standard deviation number of rings per cell in wildtype and WAVE complex-null cells; wildtype mean = 18.8 +/-27.4, n = 37 cells, WAVE complex-null mean = 3.09 +/-3.65, n = 66 cells; cells pooled from the same 3 independent experiments; unpaired t-test two-tailed ****P value <0.0001. **(E)** Images of HUVECs expressing full-length IRSp53 (left; see **S10 Video**) or I-BAR only domain (right; see **S11 Video**) at Y27632-induced TEMs. Scale bar: 5μm. **(F)** Quantification of the enrichment of full-length IRSp53 and I-BAR domain at TEMs. Violin plot of enrichment, which was measured as described in **Fig 4C**. Each time point across TEM closure was considered as a single data point: full-length n = 186 and I-BAR n = 291, both from 4 TEMs from at least 3 independent experiments per condition; Mann-Whitney test of significance two-tailed ****P value <0.0001. **(G)** Schematic summary of IRSp53 structure-function findings. Though IRSp53’s I-BAR domain is sufficient to localize to filopodia-like structures, the I-BAR domain along with its interactions with the WAVE complex are required for IRSp53 to enrich to lamellipodia, rings, and TEMs.

Since IRSp53’s SH3 domain interacts with other actin regulators besides the WAVE complex, such as Mena (60), Eps8 (62), and mDia1 (57), we tested whether IRSp53’s localization patterns are dependent on the WAVE complex by imaging full-length IRSp53-eGFP in WAVE complex depleted cells. The WAVE complex was depleted via CRISPR-Cas9-mediated knock out of Hem1, which resulted in the depletion of the other subunits (23,24,63). Upon F-actin inhibition, full-length IRSp53 failed to robustly form rings in the WAVE complex depleted cells (**Fig 6D**). These data suggest that while IRSp53’s I-BAR domain is sufficient to enrich to filopodia-like structures, IRSp53 requires its I-BAR domain and its interactions with the WAVE complex to localize to lamellipodia and saddle geometries (**Fig 6G**). Our data highlight how proteins can partner together to sense complex geometries.

To probe the generality of IRSp53 requiring both its I-BAR domain and its ability to bind to the WAVE complex to sense saddle curvature, we again used the HUVEC TEMs system, which has the advantage of providing a persistent saddle morphology. A previous study found that IRSp53’s isolated I-BAR domain fails to recognize TEMs (47), and we confirmed these results in our hands (**Fig 6E-F**). This is consistent with our observation that the isolated I-BAR construct failed to enrich to ring structures around saddle-shaped membrane invaginations in dHL60s (**Fig 6C**). Additionally, because full-length IRSp53 recognized saddle curvature in dHL60 cells (**Fig 5**), we expected that full-length IRSp53 would also recognize TEMs, and our observations are consistent with this hypothesis (**Fig 6E-F**). These data further support our finding that IRSp53 depends on its ability to interact with the WAVE complex to localize to saddle geometries (**Fig 6G**).

Though membrane invaginations during F-actin inhibition and TEMs both display saddle curvature, they sometimes differ in scale: invaginations have diameters on the nanometer scale whereas TEMs have diameters ranging from microns to fractions of a micron. Based on the characteristic size of the WAVE complex rings in the absence of actin polymer in dHL60 cells and the rings formed by IRSp53 in the presence and absence of actin polymer (**Fig 5E**), we hypothesized that this represents the optimal membrane geometry for the WAVE complex enrichment, suggesting a preference for saddles with a negative curvature of lamellipodia-size, κ ∼ 1/65 nm^-1^, and positive curvature, κ ∼ 1/135 nm^-1^. An advantage of the TEMs saddle system is that it maintains a fixed lamellipodial-like negative curvature in the z-plane while it scans a range of positive curvatures throughout its closure. Therefore, we can use this range of hole sizes to evaluate the curvature preference for the WAVE complex. Depending on the type of curvature sensor, the possible behaviors of a molecule’s local concentration (signal per unit length) during TEM closure would be that it: increases, remains constant, or decreases (**Fig 7A**). A saddle geometry sensor presented with fixed negative curvature in one axis and a range of positive curvatures in the other axis would increase its local concentration until the TEM closure reaches the sensor’s preferential positive curvature. From analyzing the WAVE complex signal throughout TEM closure, its local concentration per unit TEM perimeter increases at a rate significantly different from that of the membrane (**Fig. 7B-C**). As the imaging was performed with confocal microscopy, we are unable to resolve TEM diameters below the diffraction limit. However, the WAVE complex’s progressive enrichment to smaller and smaller TEMs toward the diffraction limit is consistent with our initial finding that the WAVE complex spontaneously forms sub-diffraction ring structures (**Fig 2B-D**). These data suggest that the WAVE complex prefers nanoscale saddle geometry in a range of cellular and physiological contexts.

**Figure 7.**
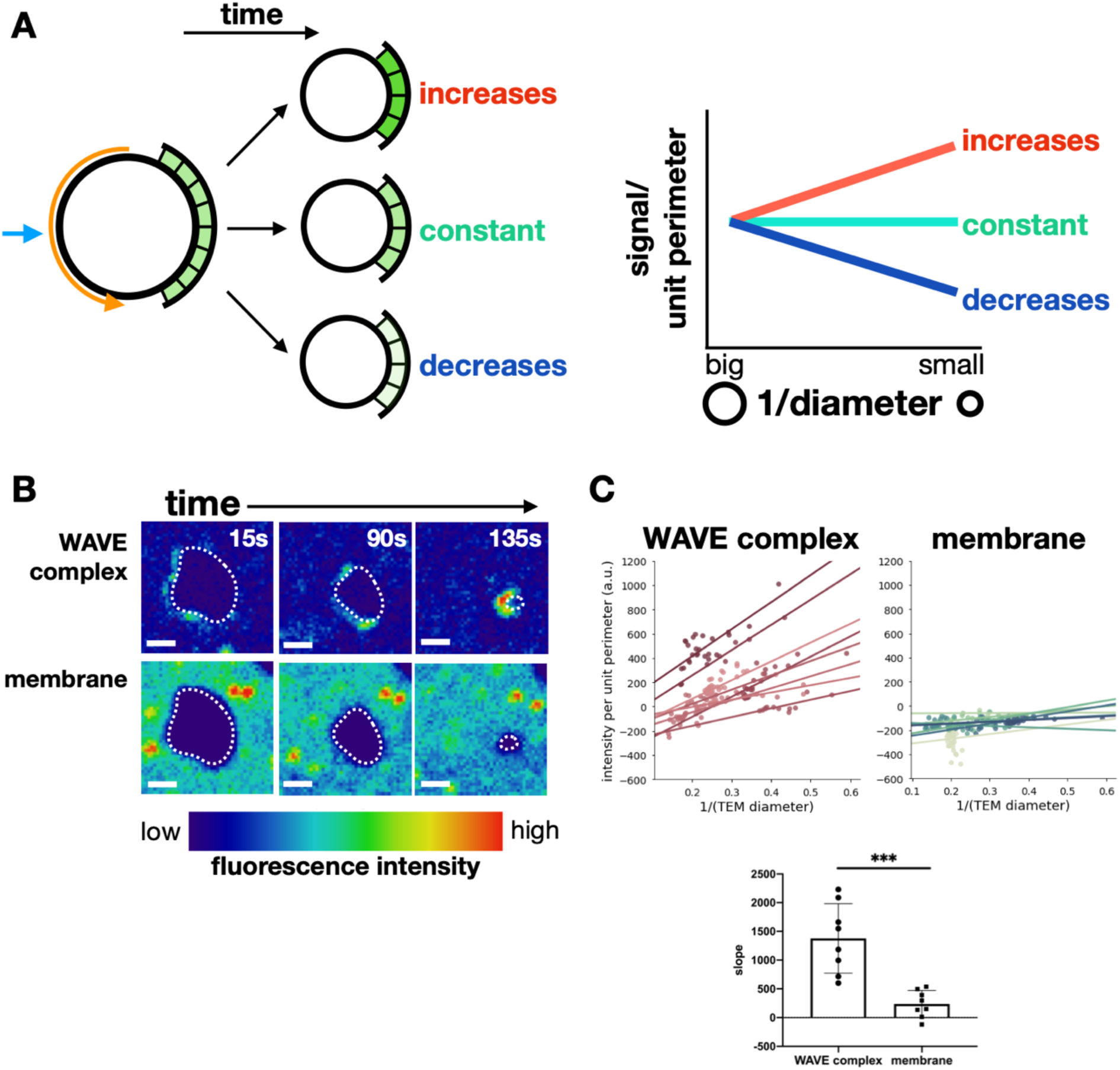
WAVE complex enriches to closing TEMs. **(A)** Schematic of different molecular behaviors during TEM closure. As TEMs close, the local concentration (signal per unit length) could increase, remain constant, or decrease. A saddle geometry sensor would increase its local concentration until the TEM closure reaches the sensor’s preferred radius of curvature. **(B)** Time-lapse images of the WAVE complex (top) and the membrane (bottom) as a TEM closes. Images in each set have the same intensity scale in a LUT that eases the visualization of signal enrichment. The dotted line outlines the TEM membrane mask. Spinning disk confocal imaging; time in seconds; scale bar: 2μm. See **S12 Video**. **(C)** Graph of the WAVE complex (top left) and membrane (top right) signal per unit perimeter as a TEM closes. Each line represents a linear regression for a single TEM over time. The WAVE complex shows higher enrichment at smaller (more positively curved) TEMs, suggesting a preference for membrane saddles with high positive curvature. Bottom: bar graph of mean +/- standard deviation of the slopes; the WAVE complex and membrane each had n = 8 TEMs from at least 3 independent experiments; unpaired t-test two-tailed ***P value <0.0002.

## DISCUSSION

### The WAVE complex assembles into a linear, curved oligomer at sites of saddle membrane curvature

In this work, we investigated how the WAVE complex achieves the linear spatial organization that is essential for lamellipodial formation. We found that the WAVE complex assembles into highly ordered, linear oligomers in the form of nanoscale rings (**Fig 2**). These rings are formed in the absence of actin polymer and associate with regions of membrane saddle curvature (**Fig 2**-**3**). The WAVE complex’s enrichment to membrane saddles is also apparent in the presence of actin polymer as it localizes to closing TEMs (**Fig 4, Fig 7**). IRSp53 requires its interactions with the WAVE complex to localize to lamellipodia, membrane invaginations, and TEMs (**Fig 5-6**). This finding highlights how proteins can partner together to sense complex geometries. The WAVE complex’s association with saddle-shaped membrane coupled with its regulation of actin polymerization likely forms the basis for emergent self-organizing behaviors like cell morphogenesis during migration and closure of transcellular holes.

How the WAVE complex achieves its oligomeric association into stereotyped 270nm rings is unknown. There is no evidence that the WAVE complex oligomerizes on its own (64). It is likely that the WAVE complex requires its interacting partners to form higher-order oligomers; this is true for N-WASP (29,31). It is possible that the same proteins that activate the nucleation ability of the WAVE complex, such as phosphoinositides, Rac, and Arf (9), are also needed to activate its oligomerization. Furthermore, our observation that the WAVE complex and IRSp53 collaborate to sense and form rings (**Fig 5-6**) suggest that IRSp53 may be a constituent of the rings.

### Parallels with septin biology for oligomeric rings as curvature sensors

The WAVE complex’s oligomerization into rings may contribute to how it recognizes sites of membrane saddle geometry. Ring formation has been observed for other proteins capable of sensing membrane curvature. Septins form rings upon treatment of actin destabilizing drugs *in vivo* and purified septins form similarly-sized rings *in vitro* (65). The radius of curvature of septin rings reflects septins’ intrinsic curvature preference both *in vitro* and *in vivo* (65,66). Like septins, the WAVE complex also forms ring structures upon actin depolymerization, and these rings associate with curved membranes (in the case of the WAVE complex sites of saddle curvature with a radius of positive curvature of 135nm). Thus, the geometry of the WAVE complex’s nanoscale rings may predict the WAVE complex’s preferred geometry. Consistent with this idea, the WAVE complex prefers membrane saddles with near-diffraction-limited curvature in the positive axis for closing transendothelial cell macroaperture tunnels (**Fig 7B-C**) and overexpression of IRSp53 generates similarly sized rings even in the presence of an intact actin cytoskeleton (**Fig 5C-E**).

What is the dimension of the WAVE complex rings in the negative curvature axis? Although we cannot visualize this directly in our TIRF-SIM images of F-actin inhibited cells (negative curvature axis is in the z plane where TIRF provides no information), our electron microscopy imaging of membrane invaginations gives a height estimate of ∼130nm, which is consistent with the negative curvature of the edge of lamellipodia in our work (**S4D Fig**) and others’ work (67–69), and consistent with the ∼100nm resolution of TIRF-SIM imaging of propagating WAVE complex waves in non-latrunculin treated cells. The presence of IRSp53 in the rings and its partnering interactions with the WAVE complex would suggest the negative curvature axis is set by IRSp53, which has an intrinsic curvature of around 20nm in the negative axis and curvature sensing up to around 100nm (70). If protein oligomerization is setting the positive axis of curvature sensing, this could account for why the WAVE complex can tolerate a range of positive curvatures: from no curvature/flat (some portions of lamellipodia) across many microns (in large TEMs) up to the preferred curvature of diffraction-limited TEMs and nanoscale membrane invaginations (**Fig 2, 7**).

### IRSp53 partners with the WAVE complex to recognize saddle geometry

How does the WAVE complex recognize saddles? Saddle enrichment could be accomplished with one protein sensing the positive curvature and another sensing the negative curvature. The WAVE complex is not known to sense curvature on its own, but it interacts with IRSp53, a member of the nanoscale negative curvature sensing BAR domain family. By comparing full-length IRSp53 to its I-BAR only domain in dHL60s, HUVECs, and in WAVE complex depleted dHL60s, we clarified how IRSp53’s localization is guided by its partnering interactions with the WAVE complex to enrich to lamellipodia and sites of saddle curvature (**Fig 6**). Others have also observed differences between full-length IRSp53 and its I-BAR only domain with super-resolution imaging. For example, with STORM imaging of filopodia in mouse neuroblastoma cells, IRSp53 localizes to the lateral edges of filopodia, whereas its isolated I-BAR domain is uniformly distributed in filopodia (61). Our work highlights how proteins partner together for complex, saddle shape sensing. What about the WAVE complex and IRSp53 interaction enables IRSp53 to localize to lamellipodia and rings? The oligomeric arrangement of IRSp53 on its own may be different than that seen in conjunction with the WAVE complex. This merits future investigation.

### Saddle curvature recognition of the WAVE complex may result in emergent behaviors, such as self-straightening lamellipodia and sealing of transcellular holes

How are the WAVE complex’s nanoscale rings related to its excitable dynamics during neutrophil migration? We interpret the WAVE complex rings that associate with saddle curvature to represent its preferred geometry of oligomerization, with a 135nm radius of curvature in the positive axis and around 65nm radius of curvature in the negative axis.

The coupling of saddle curvature recognition (cell shape) and patterning F-actin polymerization (physical forces) could result in a number of emergent behaviors. For example, if the WAVE complex both recognizes and generates sites of membrane saddle curvature, this could form a feedback loop that organizes the expansion and self-straightening of lamellipodia. Near the initiation of a lamellipodium, a small sheet-like membrane deformation has saddle curvature at its lateral edges (**Fig 8**). WAVE complex association with these saddles would result in a laterally expanding zone of actin nucleation, where the wave would grow at the sides but be confined to a small region at the tip of the lamellipodium based on the negative curvature sensing. This spatial constraint on the positive feedback arm of the excitable actin nucleation circuit could explain why the domain of activation for this circuit is much thinner than other excitable circuits, such as cortical excitability in *Xenopus* (71) and Min protein oscillations for bacterial cell division (72).

**Figure 8.**
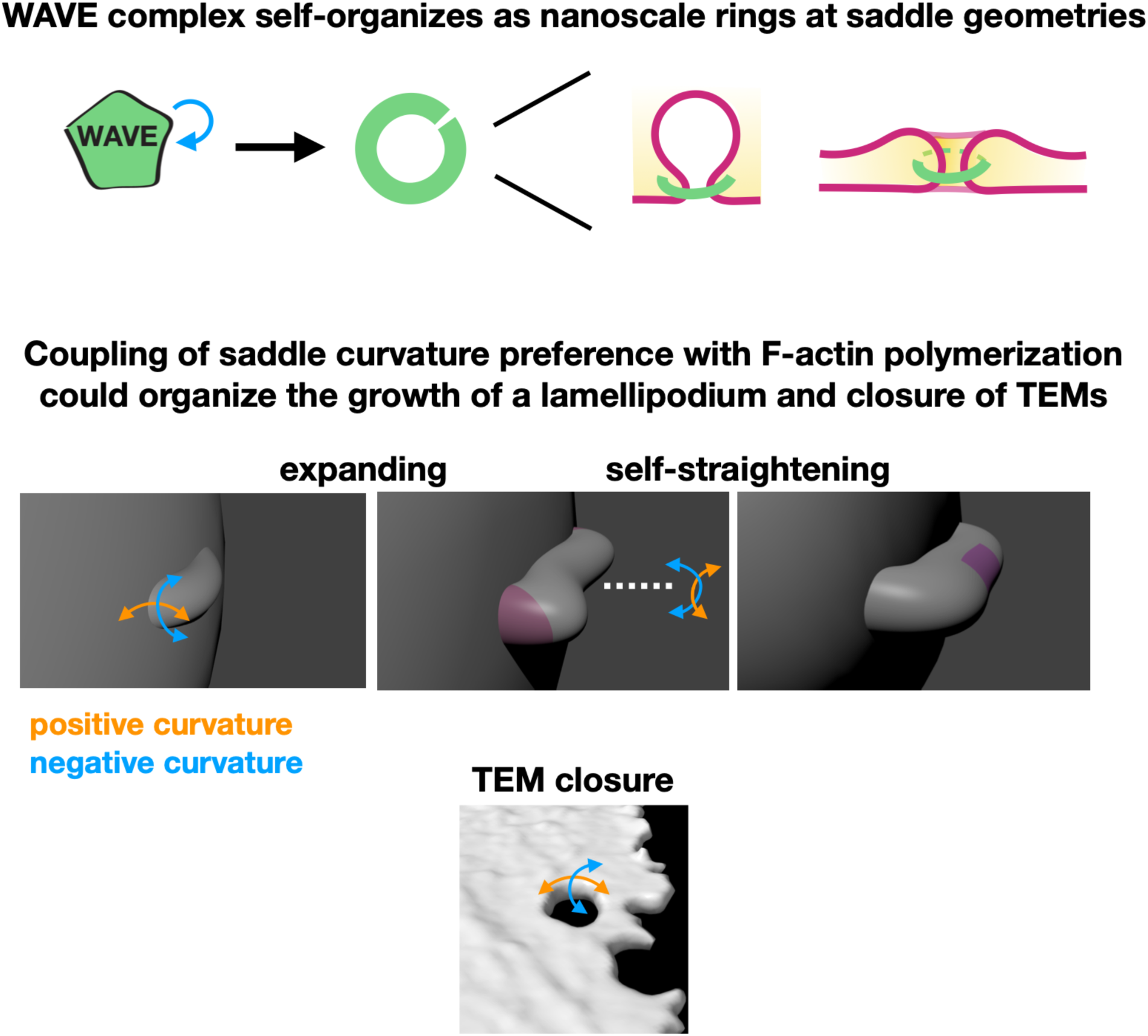
Summary and model. **(A)** The WAVE complex self-organizes as linear, oligomeric structures at saddle membrane geometries. **(B)** The coupling of saddle curvature enrichment with F-actin polymerization could organize the growth of a lamellipodium and closure of TEMs. At the initiation of a lamellipodium, the lateral edges exhibit saddle curvature. WAVE complex association would lead to the growth of the extension (middle panel highlights extended membrane). Portions that “lag” behind also exhibit saddle curvature. WAVE complex association would result in self-straightening behavior (right panel highlights straightened membrane). Furthermore, WAVE complex association at the saddle geometry of TEMs would lead to TEM closure (bottom panel).

Saddle enrichment would also enable maintenance of a uniform leading edge advancement — any regions of the lamellipodia that lag behind also display saddle curvature, resulting in flattening of the front (**Fig 3D-E, Fig 8**). If the lamellipodium requires a negatively curved edge, the flattening of this structure as it contacts a barrier could provide a geometric mechanism of barrier avoidance; the WAVE complex is known to extinguish at mechanical barriers (7). Finally, saddle recruitment of the WAVE complex could enable the recognition and actin-based sealing of transient endothelial macroapertures (47). Simple local rules of protein association could generate a number of the emergent features of cell shape and movement.

## MATERIALS AND METHODS

### Cell culture

All cells were cultured in a 37°C/5% CO_2_ incubator.

*HL60s* - HL60 cells were cultured in RPMI-1640 with 25mM HEPES (Corning) with 10% (vol/vol) heat-inactivated fetal bovine serum (FBS; Gibco) and maintained at 0.2-1.0 × 10^6^ cells/mL. Cells were differentiated with 1.5% DMSO (Sigma-Aldrich) in growth media for 4-5 days prior to experiments. All imaging was done with differentiated HL60s unless otherwise stated. Hem1-depleted HL60 cells were from (24).

*HEK-293Ts* - HEK-293T cells were used to generate lentivirus. 293Ts were cultured in DMEM (Gibco) with 10% FBS. Cells were cultured up to 80% confluency.

*HUVECs* - HUVEC cells were a kind gift from the MS Conte lab (University of California, San Francisco [UCSF]). HUVECs were cultured in HyClone M-199 media (GE Healthcare) with 1% antibiotics-antimicrobial (Gibco), 15.3 units of heparin (Sigma), 1 vial of endothelial cell growth supplement (EMD Millipore), and 10% FBS. Cells were cultured up to 80% confluency and growth media was replaced every other day. Experiments used cells with fewer than 10 passages.

*B16F10* - B16F10 cells were cultured in DMEM (Gibco) with 10% FBS, 1% penicillin-streptomycin (Gibco), and 1% GlutaMax (Life Technologies). Cells were cultured up to 80% confluency.

*U2OS* - U2OS cells were a kind gift from the M von Zastrow lab (UCSF). U2OS cells were cultured in McCoy’s 5A media (Gibco) with 10% FBS. Cells were cultured up to 80% confluency.

### Plasmids

For stable expression, all constructs were cloned into the pHR lentiviral backbone (provided by the RD Vale lab, UCSF) with a SFFV promoter via standard Gibson assembly. The other WAVE complex subunit constructs (eGFP-Abi1, eGFP-Abi2, eGFP-Sra1, eGFP-Nap1, and eGFP-WAVE2) were a kind gift from the G Scita lab (European Institute of Oncology). IRSp53 was from the DNASU Plasmid Repository (pENTR223) and subsequently subcloned into a pHR vector. IRSp53^1-250^ was “I-BAR,” the N-terminal 250 amino acids, and IRSp53^251-512^ was “C-term,” the C-terminal 262 amino acids beyond the I-BAR domain. For all IRSp53 constructs, the eGFP was C-terminally tagged.

### Cell line generation

#### Lentivirus

All HL60s and some HUVEC lines were stably expressing the constructs of interest. HEK-293Ts were plated in 6-well plates (Corning) until 70-80% confluency. Cells were transfected with 1.5µg of the pHR plasmid along with two plasmids containing the lentiviral packaging proteins (0.167µg of pMD2.G and 1.3µg of p8.91) with Trans*IT*-293 (Mirus Bio). After 2-3 days of transfection, lentivirus-containing supernatant was collected, filtered with a 0.45µm filter (EMD Millipore), and concentrated 20x with Lenti-X Concentrator (Takara). Lentivirus was used immediately or kept at −80°C. For HL60 transduction, 3.2×10^5^ cells, 4µg/mL polybrene, and ∼130µL of concentrated virus were incubated overnight. Cells were FACS sorted (BD Aria2). For eGFP-tagged IRSp53 structure-function constructs, cells were sorted for comparable fluorescence expression levels (**S6 Fig**). For HUVEC transduction, a 50% confluent 6-well was incubated with 4µg/mL polybrene, and ∼130µL of 1x virus overnight.

#### Electroporation

For transient expression of plasmids in HUVEC cells, electroporation was used. For each electroporation condition, 50,000 cells were resuspended in 10µL of Buffer R (Thermo Scientific) with 500ng of DNA. Electroporations were performed with the Neon Transfection System (ThermoFisher) with 10µL tips at 1350V for 30ms for 1 pulse. Cells recovered in 500µL of culture media and imaged the next day.

#### Lipofectamine

For B16F10 and U2OS cell lines, plasmids were transiently expressed with Lipofectamine 2000 or 3000 (Invitrogen) per manufacturer’s instructions.

### Imaging

#### Cell preparation

##### HL60s

For imaging, differentiated cells were resuspended in imaging media (either Leibovitz’s L-15 [Gibco] with 0.5% FBS or mHBSS [10x stock consists of 1500mM NaCl, 40mM KCl, 10mM MgCl_2_, 12mM CaCl_2_, 100mM glucose, 200mM HEPES, pH 7.2]). Cells were plated onto fibernectin coated wells (100 µg/mL for 1hr at room temperature) and incubated (37°C/5% CO_2_) for at least 7 minutes before 2-3 washes with imaging media. For chemoattractant stimulation, a 2x stock of 20nM fMLP (Sigma) was added. For additional chemoattractant stimulation, a 2x stock of 200nM fMLP was added. For F-actin inhibition, a 2x stock of 1µM latrunculin B (EMD Millipore and Sigma) with 200nM phorbol 12-myristate 13-acetate (PMA; Sigma; for persistent Hem1 activation) was used, unless noted otherwise. Other F-actin inhibition drugs were latrunculin A (EMD Millipore) and cytochalasin B (Sigma). All initial stocks were dissolved in 100% dry DMSO and freshly diluted in imaging media before experiments.

##### HUVECs

To induce TEM formation, HUVECs were incubated in 50µM Y27632 (Tocris and Sigma) for at least 4 hours before imaging. Y27632 was present throughout imaging.

##### Membrane labeling

For membrane labeling, Vybrant DiD (Invitrogen) or CellMask DeepRed (Invitrogen) was freshly diluted (0.5-1X) in imaging media. HL60s were labeled in suspension for 30sec at 37°C and washed 2-3 times with imaging media. Adherent cell lines were labeled for 5min at 37°C and washed 3-5 times with phosphate buffered saline (PBS).

##### Fixation

Cells were plated in #1.5, 8-well Lab-Tek II chambers (Thermo Fisher Scientific). To fix cells, media was aspirated while simultaneously adding 200µL of 2% glutaraldehyde (Sigma-Aldrich) for 10min and washed with PBS for 30sec and 60sec. Aldehydes were quenched with 0.1% sodium borohydride (Sigma-Aldrich) for 7min and followed by 2-3 10min PBS washes. Samples were imaged immediately or stored at 4°C. All dilutions were prepared fresh in cytoskeleton buffer with sucrose (CBS), a TJ Mitchison lab (Harvard Medical School) recipe: 10mM MES pH 6.1, 138mM KCl, 3mM MgCl, 2mM EGTA, with 0.32M sucrose added fresh before use. HUVECs were fixed with 4% PFA in PBS, pH 7.4 instead of with 2% glutaraldehyde. All materials were warmed to 37°C.

#### Microscopes

##### TIRF-SIM

TIRF-SIM imaging was performed on DeltaVision OMX SR (GE Healthcare) with a 60x/1.42 numerical aperture (NA) oil Plan Apo objective (Olympus) with 1.516 or 1.518 refractive index oil (Cargille). HL60 experiments were performed at room temperature while adherent cells were performed at 37°C/5% CO_2_. Images were processed with softWoRx (OMX SI Reconstruction with the default parameters including Wiener filter: 0.001; OMX Align Images with the Legacy Image Registration method). Display of TIRF-SIM membrane images were smoothed in Fiji.

##### Spinning disk confocal

Images were acquired on a Nikon Eclipse Ti microscope with a 60x/1.40 NA Plan Apo objective (Nikon), Yokogawa CSU-X1 spinning disk confocal, and a Prime 95B cMOS camera (Photometrics). 405, 488, 561, 640nm laser lines (Agilent Technologies) and environmental control (37°C, 5% CO_2_; Okolab) was used. Software was controlled with Nikon Elements.

##### Lattice light sheet and processing

Lattice-light sheet imaging was performed in a manner previously described (73) and followed an established protocol (74). Briefly, 5mm round coverslips (Warner Instruments) were cleaned by plasma cleaning and coated with fibernectin (100 µg/mL for 1hr). Cells were plated as described above. The coverslip sample was then loaded into a sample holder and placed into the previously conditioned microscope sample bath (with 25nM chemoattractant) and secured. Imaging was performed with a 488nm laser (MPBC). Camera exposure was 10ms per frame leading to a temporal resolution of 2.25 sec in single color mode.

Raw image files were deconvolved using the iterative Richardson-Lucy algorithm with the known point spread function for each channel, which were collected prior to each day of imaging (73). The code for this process was provided by the E Betzig lab (Janelia Research Campus) originally written in Matlab (The Mathworks) and ported into CUDA (Nvidia) for parallel processing on the graphics processing unit (GPU, Nvidia GeForce GTX Titan X). Each sample area underwent 15 iterations of deconvolution.

Regions of interest within the sampling were cropped down to size and compressed from 32-bit TIFFs to 16-bit TIFFs using in-house Matlab code to allow immigration into the 3D visualization software ChimeraX (UCSF Resource for Biocomputing, Visualization, and Informatics (75)). To highlight intensity ranges, additional channels were created by thresholding.

##### Transmission electron microscopy

Cells were prepared for convention EM in two ways: cells were either plated on fibernectin-coated ACLAR discs (TedPella), fixed with glutaraldehyde as described above and dehydrated or cells were plated on sapphire disks and fixed via high pressure freezing/freeze-substitution. For the glutaraldehyde fixation method, cells were treated with 100nM chemoattractant or 500nM latrunculin B for 5min, fixed, stained with uranyl acetate and OsO_4_, dehydrated with cold ethanol, and embedded with Epon 812 resin. For the high pressure freezing/freeze-substitution method, cells were plated onto 3mm diameter sapphire disks and treated with 100nM chemoattractant or 500nM latrunculin B for 5min prior to freezing. The sapphire disk was then placed, cells toward the inside of a 100µm well specimen carriers (Type A, Technotrade International Inc) containing 20% bovine serum albumin (BSA) in growth media. The sandwiched cells were frozen using a BalTec HPM 01 high pressure freezer (BalTec). Freeze-substitution in 1% OsO_4_, 0.1% uranyl acetate, 1% methanol in acetone, containing 3% water (76,77) was carried out with a Leica AFS2 unit. Following substitution, samples were rinsed in acetone, infiltrated and then polymerized in Eponate 12 resin (Ted Pella). For conventional electron microscopy, serial 50nm sections were cut with a Leica UCT ultramicrotome using a Diatome diamond knife, picked up on Pioloform coated slot grids and stained with uranyl acetate and Sato’s lead (78). Sections were imaged with a FEI Tecnai T12 TEM at 120 kV using a Gatan 2k × 2k camera. For EM tomography, 200nm sections were cut, mounted on grids and stained as above for serial thin sections. Tomograms were acquired using an FEI T20 TEM at 200 kV, and tomograms reconstructed using eTomo/IMOD (79,80).

### Image analysis and statistics

All image analysis was performed in Fiji and/or Python. Data handling and statistical tests were performed with Python and Prism 8 (GraphPad).

#### Diameter calculation

After background subtraction (rolling ball), segmentation was performed with Trainable Weka Segmentation (81) machine learning algorithm to identify ring structures. The diameter of a perfect circle with the same perimeter of the ring particle was calculated (diameter = C/*π*; C=circumference).

#### Membrane curvature and velocity

Fluorescence images were segmented using a 3-step process consisting of Gaussian smoothing, intensity-based thresholding, and binary erosion. The threshold and degree of erosion were chosen manually to align the boundary of the binary image with the apparent edge of the cell membrane. To facilitate temporal analysis of edge properties, these boundaries were then fit using a spline interpolation consisting of 1,000 points. Edge velocity at a particular point, P, at time, t, was estimated by calculating the average of the distance transforms of the binary images at times t-1 and t+1 and interpolating the value of this function at the coordinates of P. The radius of curvature at a point, P, was calculated by approximating the radius of the osculating circle, C, along the boundary at the coordinates of P. To approximate this osculating circle, we chose a scale parameter, S, then collected the coordinates of the two points at indices S units away from P. Given these three points, a unique equation of a circle passing through them was calculated. A sign was assigned by comparing the vector between P and the center of circle C and the normal vector of the boundary.

#### EM curvature

After identifying the plasma membrane, a perfect circle was fitted to both sides of the neck of an invagination and at the tip of a protrusion. Curvature, *κ*, was defined as *κ* = 1/R (R = radius).

#### IRSp53 and Hem1 colocalization

Images were background subtracted (rolling ball) and Fiji’s coloc2 function was applied with the segmented images and an ROI of the combined masks.

#### IRSp53 lamellipodia/filopodia enrichment

After background subtraction (rolling ball), two masks were generated: a lamellipodia mask was created from Hem1 signal and a filopodia mask was created from the eroded regions of a cell mask. After subtracting the cytosolic background from each segment (cytosolic intensity per unit area multiplied by the segment’s area), the intensity per unit area of each segment was calculated, i.e. lamellipodia signal/lamellipodia area. The ratio of the lamellipodia to the filopodia intensities per unit areas was graphed.

#### IRSp53 structure function ring enrichment

After background subtraction (rolling ball), a ring mask was generated from segmenting Hem1 signal and a background mask was generated by dilating the ring mask. IRSp53 signal was measured from both masks. After background correction (subtracting the background intensity per unit area multiplied by the ring mask area), the enrichment factor was calculated as the ratio of the ring to the background intensities per unit areas. Enrichment of all rings within a cell was averaged and treated as an individual data point.

#### TEMs enrichment

TEMs were identified by segmenting the membrane channel (TEM mask). The signal mask was generated by dilating the TEM mask by 2-3 pixels and the background mask was generated by dilating the signal mask by 2-3 pixels. A doughnut-shaped signal segment was calculated by subtracting the TEM mask from the signal mask and a doughnut-shaped background segment was calculated by subtracting the signal mask from the background mask. The doughnut-shaped signal segment was background corrected (subtracting the background intensity per unit area multiplied by the signal mask area). The enrichment factor was measured as the ratio of the signal to background intensities per unit areas. The intensity per unit perimeter was calculated as the background-corrected signal intensity divided by the signal perimeter.

## ACKNOWLEDGEMENTS

We thank the members of the Weiner lab for many conversations and support throughout the project. We want to acknowledge Wallace Marshall, Sophie Dumont, Ron Vale, Adam Frost, and Nir Gov for their helpful discussions. We thank Jessica Sherry, Kirstin Meyer, and Brian Graziano for critical reading of the manuscript. We thank DeLaine Larsen and Kari Herrington of the UCSF Imaging Core as well as Chris Rieken, Galo Garcia, David Castaneda-Castellanos, and Yina Wang for their microscopy expertise. We also thank Richard Fetter and the staff at the University of California Berkeley Electron Microscope Laboratory for advice and assistance in electron microscopy sample preparation and data collection. This work was supported by an AHA Predoctoral Fellowship (AP), NIH F31 HL143882 (RMB), NSF Predoctoral Fellowships (JPT, NTT), NIH GM118167 (ODW), the NSF Center for Cellular Construction (DBI-1548297), and a Novo Nordisk Foundation grant for the Center for Geometrically Engineered Cellular Systems (NNF17OC0028176). The UCSF Imaging Core was supported by the Research Evaluation and Allocation Committee (REAC), the Gross Fund, and the Heart Anonymous Fund. The FACS work was supported in part by the HDFCCC Laboratory for Cell Analysis Shared Resource Facility through a grant from the NIH (P30CA082103).

---

**S1 Figure.**
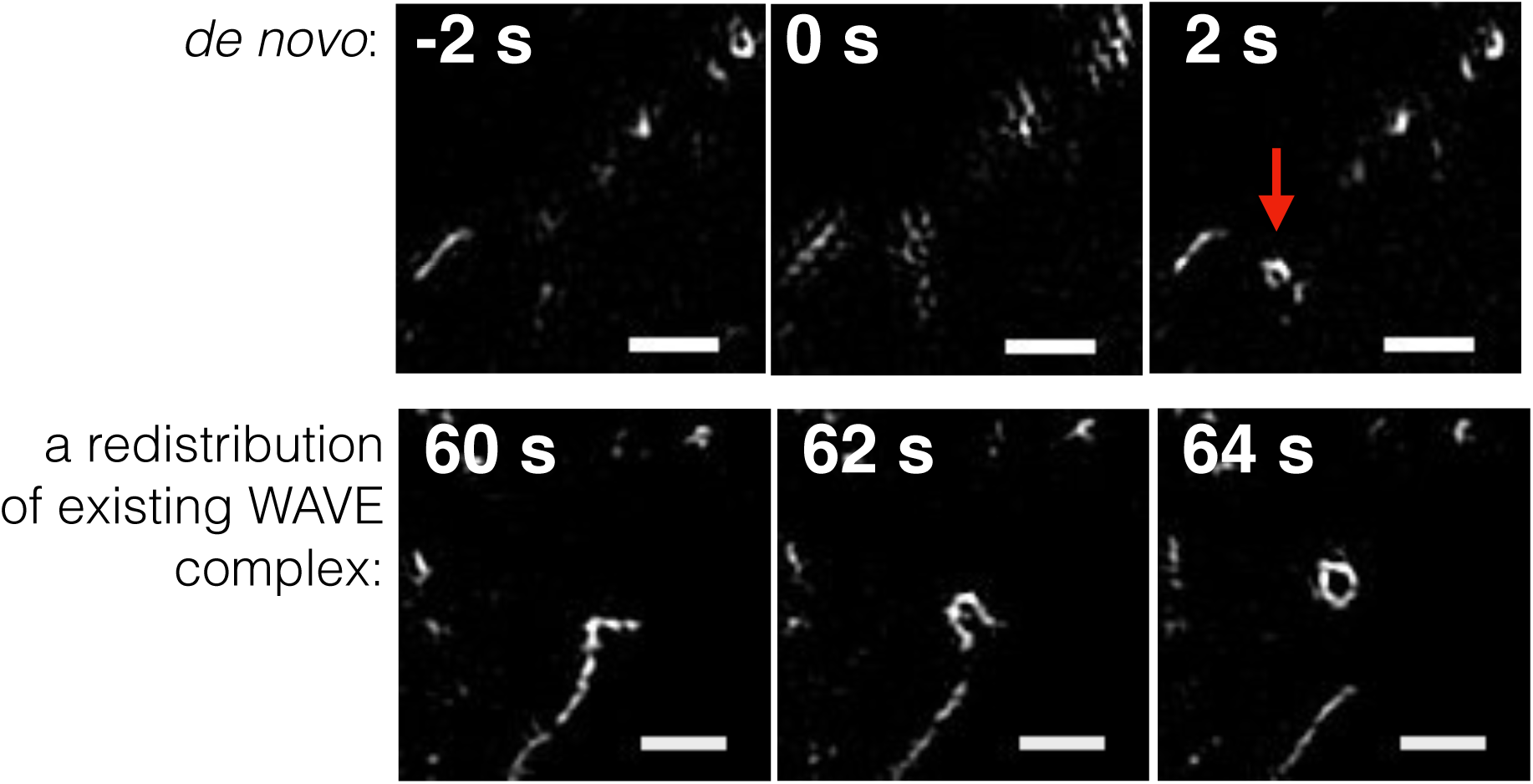
Different modes of the WAVE complex ring formation. WAVE complex (Hem1-eGFP) rings form either: *de novo* by the recruitment of new WAVE complex to the plasma membrane upon additional chemoattractant stimulation (100nM fMLP) in latrunculin B-treated cells (top) or a redistribution of existing membrane-bound WAVE complex that locally “collapses” into rings (bottom). Time T = 0 refers either to acute stimulation of either chemoattractant to latrunculin B-treated cells (top) or latrunculin B (bottom). Scale bar: 500nm.

**S2 Figure.**
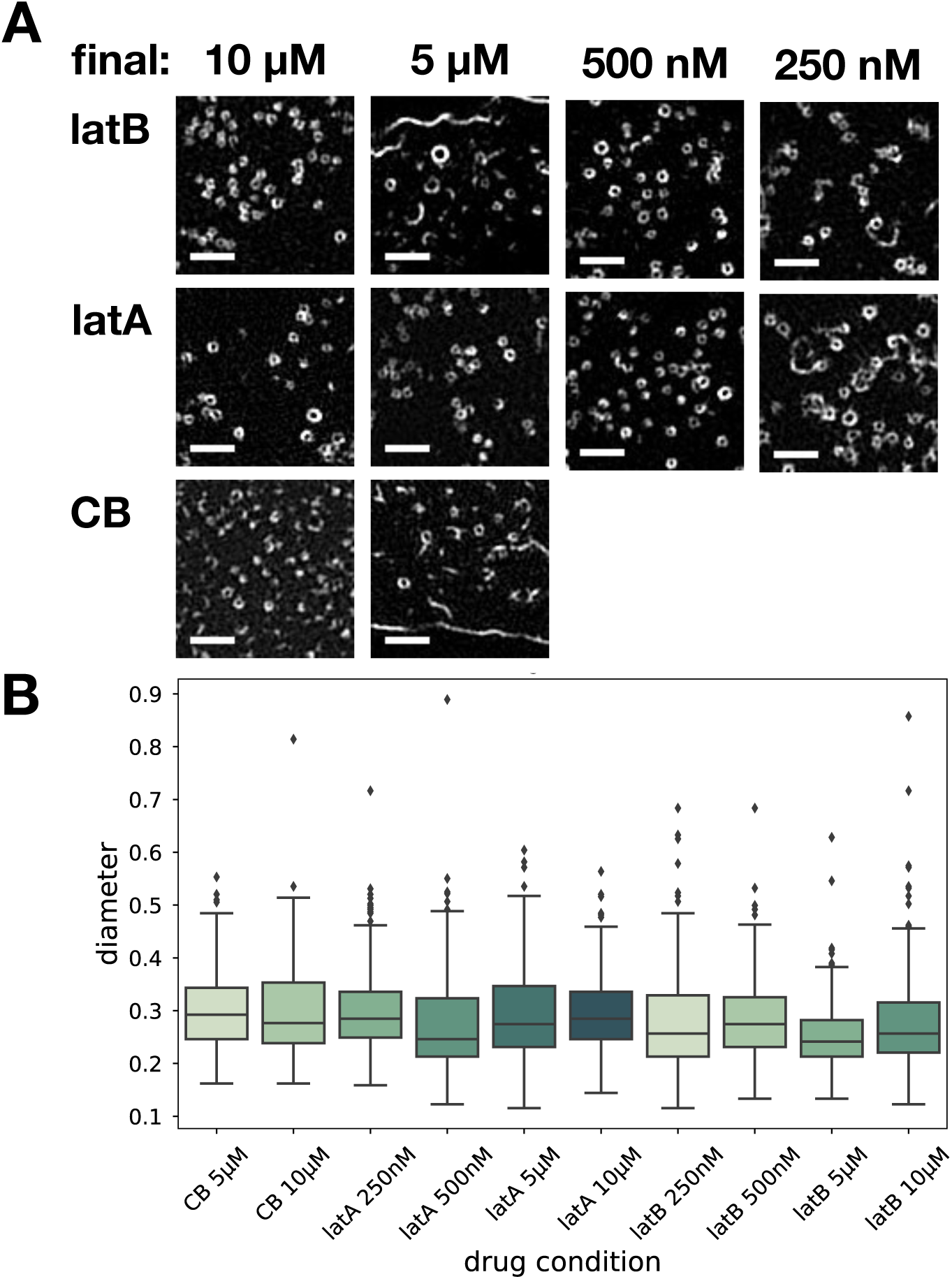
WAVE complex forms invariant ring structures across different F-actin inhibitors and concentrations. (**A**) Zoomed-in TIRF-SIM images of Hem1-eGFP rings across different drug conditions: 250nM, 500nM, 5μM, 10μM (final) of latrunculin B (latB), latrunculin A (latA), and cytochalasin B (CB). Scale bar: 1μm. **(B)** Boxplots (of interquartile range) of ring diameters across drug conditions as measured in **Fig 2D**. Each condition has at least 10 cells from at least 3 independent experiments per condition; ensemble histogram in **Fig 2D**.

**S3 Figure.**
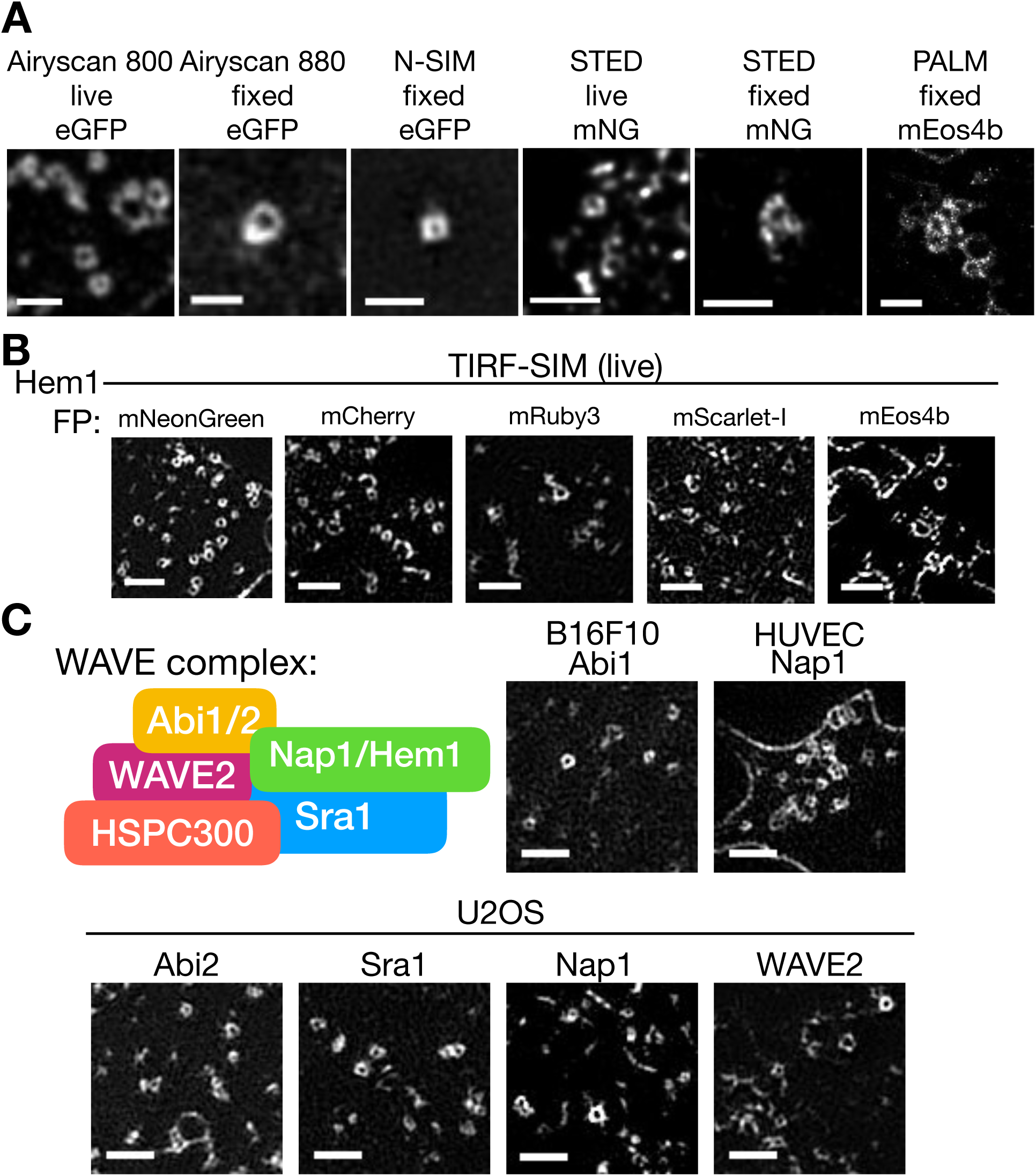
WAVE complex ring structures are observed independent of super-resolution modality, tagged fluorophore, cell type, and specific subunit. **(A)** Multiple super-resolution techniques in live or fixed cell conditions of Hem1 tagged with different fluorescent proteins all show ring structures. Microscopes and techniques used: Airyscan 800 (Zeiss), Airyscan 880 (Zeiss), Nikon-SIM (N-SIM), STimulated Emission Depletion (STED) on Leica SP8 with mNeonGreen [mNG], and photoactivated localization microscopy (PALM) on B Huang lab (UCSF) microscope. All cells treated with latrunculin B (500nM). Scale bar: 1μm. **(B)** Hem1 tagged with different fluorescent proteins show ring structures. All imaged with TIRF-SIM and treated with latrunculin B (500nM). Scale bar: 1μm. **(C)** Different WAVE complex subunits in other cell lines show ring structures. Top left, cartoon of the WAVE complex subunits. Different eGFP tagged WAVE complex subunits in B16F10 *(Mus musculus* skin melanoma cells), HUVECs (human umbilical vein endothelial cells), and U2OS (*Homo sapiens* bone osteosarcoma) cell lines. All cells treated with latrunculin B (500nM). Scale bar: 1μm.

**S4 Figure.**
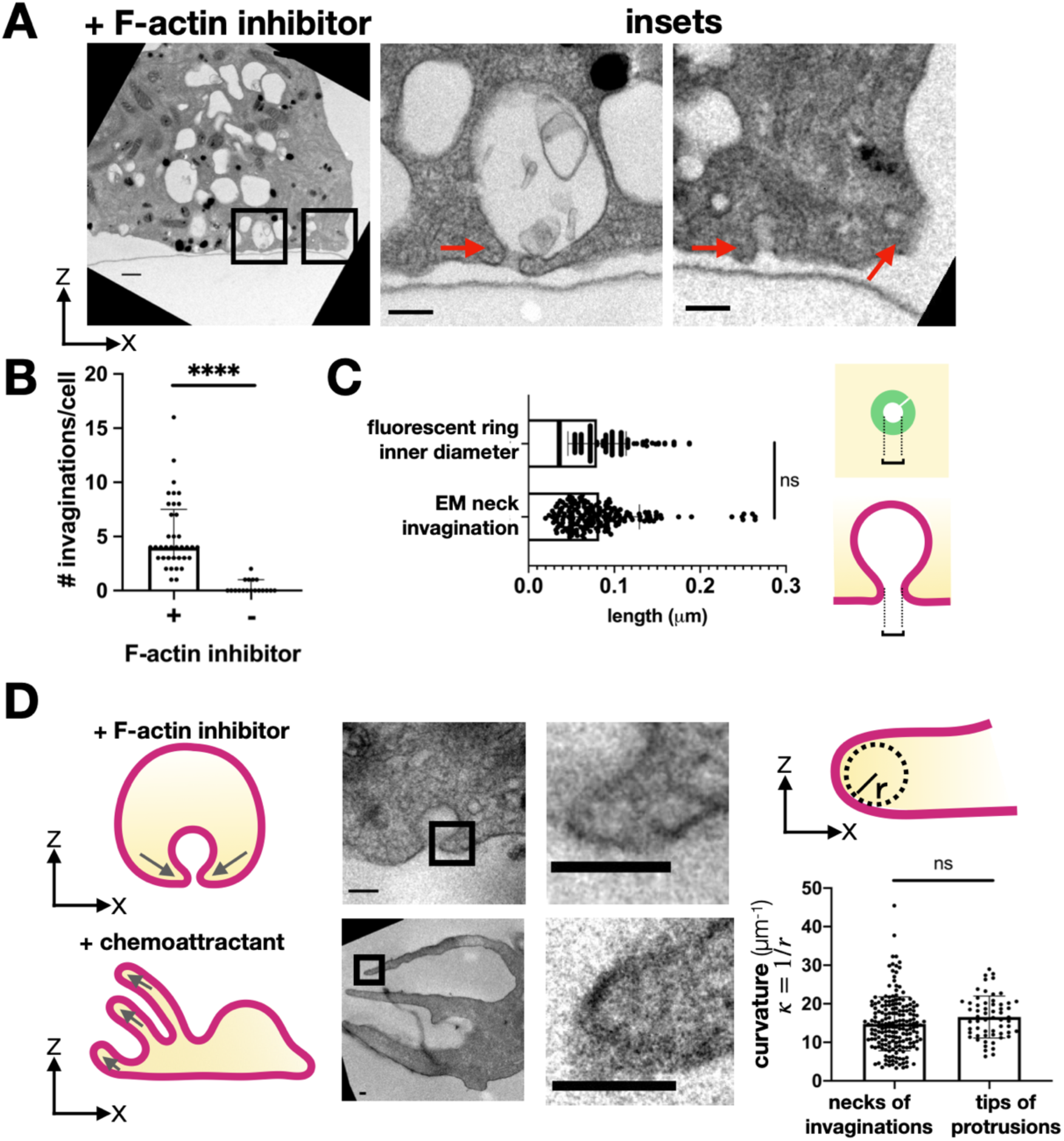
Membrane invaginations are observed by electron microscopy (EM) **(A)** Transmission electron micrographs of a cross-section of dHL60s treated with latrunculin B (500nM). Center and right micrographs are insets (black boxes on left micrograph). Red arrows point to membrane invaginations. Scale bars: 500nm (left) and 200nm (inset). **(B)** Graph comparing the number of membrane invaginations per serial section of cells treated with or without F-actin inhibitor (500nM latrunculin B). Graph shows median with interquartile range; (+) F-actin inhibitor n = 38 cells, (-) F-actin inhibitor n = 18 cells; Mann-Whitney test with two-tailed ****P value <0.0001. **(C)** Graph comparing the length across the neck of invaginations by EM to the inner diameter of the fluorescent Hem1 rings of dHL60s treated with latrunculin B (500nM). Schematic represents where measurements were made. Graph shows median with interquartile range; EM of neck invaginations n = 190 (from 38 cells), fluorescent ring inner diameter n = 222 (from 4 cells); Mann-Whitney test with two-tailed P value ∼0.3881. This finding supports the possibility that the WAVE complex enriches around the necks of membrane invaginations. Note that the invaginations have various sizes (**A**), yet the length across the neck is consistently ∼71nm. **(D)** Curvature comparison of the necks of invaginations and tips of protrusions. Electron micrographs of dHL60s treated with F-actin inhibitor (top; 500nM latrunculin B) and dHL60s treated with chemoattractant (bottom; 100nM fMLP) show that the curvature at the necks of invaginations and the tips of protrusions are similar. Graph shows mean +/-standard deviation; invagination mean = 14.68 +/-6.97μm^-1^, n = 190 (from 28 cells), protrusions mean = 16.56 +/-5.44μm^-1^, n = 59 (from 19 cells); unpaired t-test two-tailed ns P value > 0.05. Curvature, κ, defined as 1/r where r is the radius. Scale bar: 100nm.

**S5 Figure.**
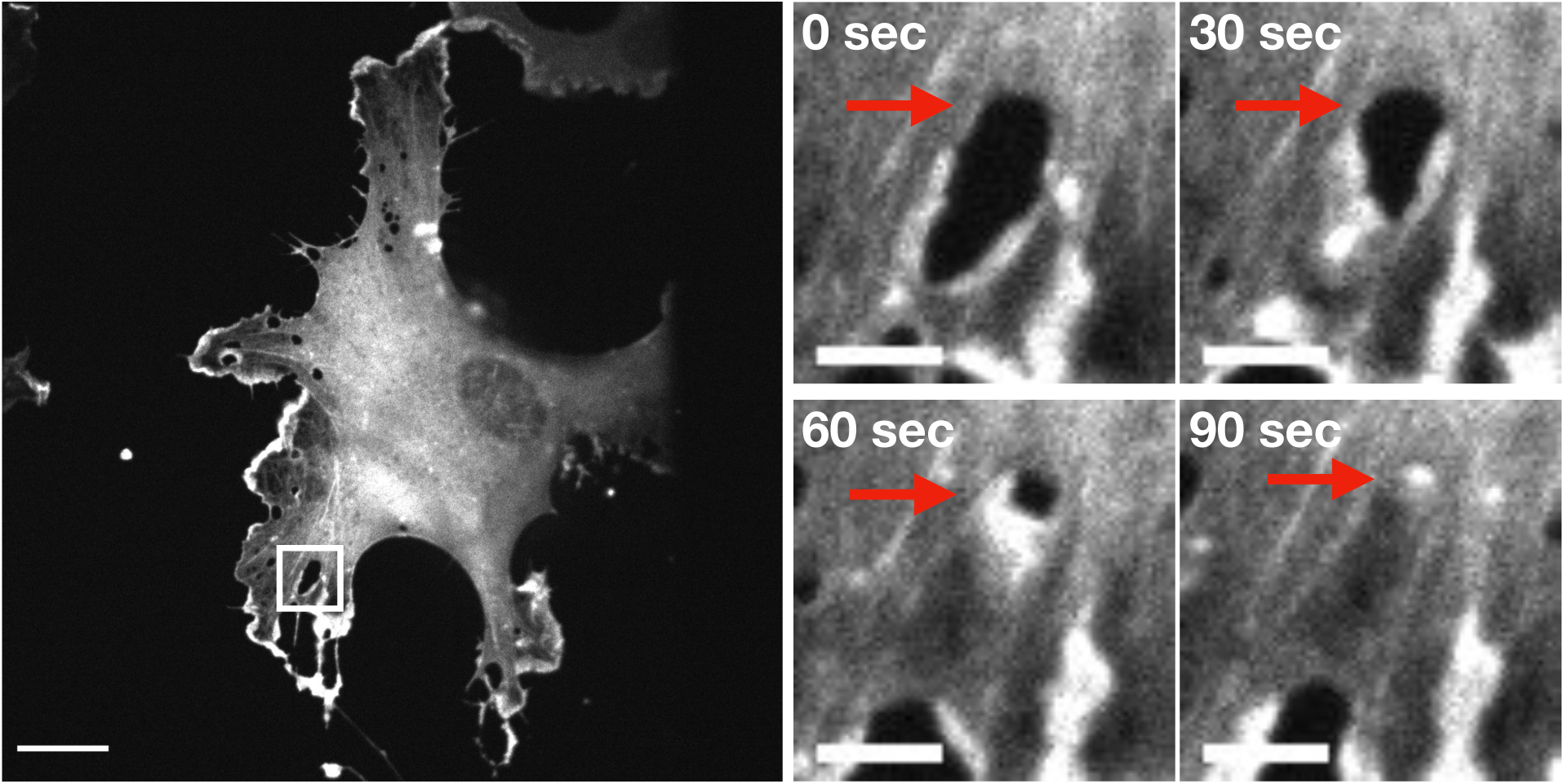
Actin localization at TEMs. HUVEC cell expressing actin-eGFP and treated with ROCK inhibitor Y27632 (50μM). Right insets: TEM closure over time; red arrows point to a TEM. Spinning disk confocal imaging; time in seconds; scale bars: 20μm (left) and 5μm (insets). See **S7 Video.**

**S6 Figure.**
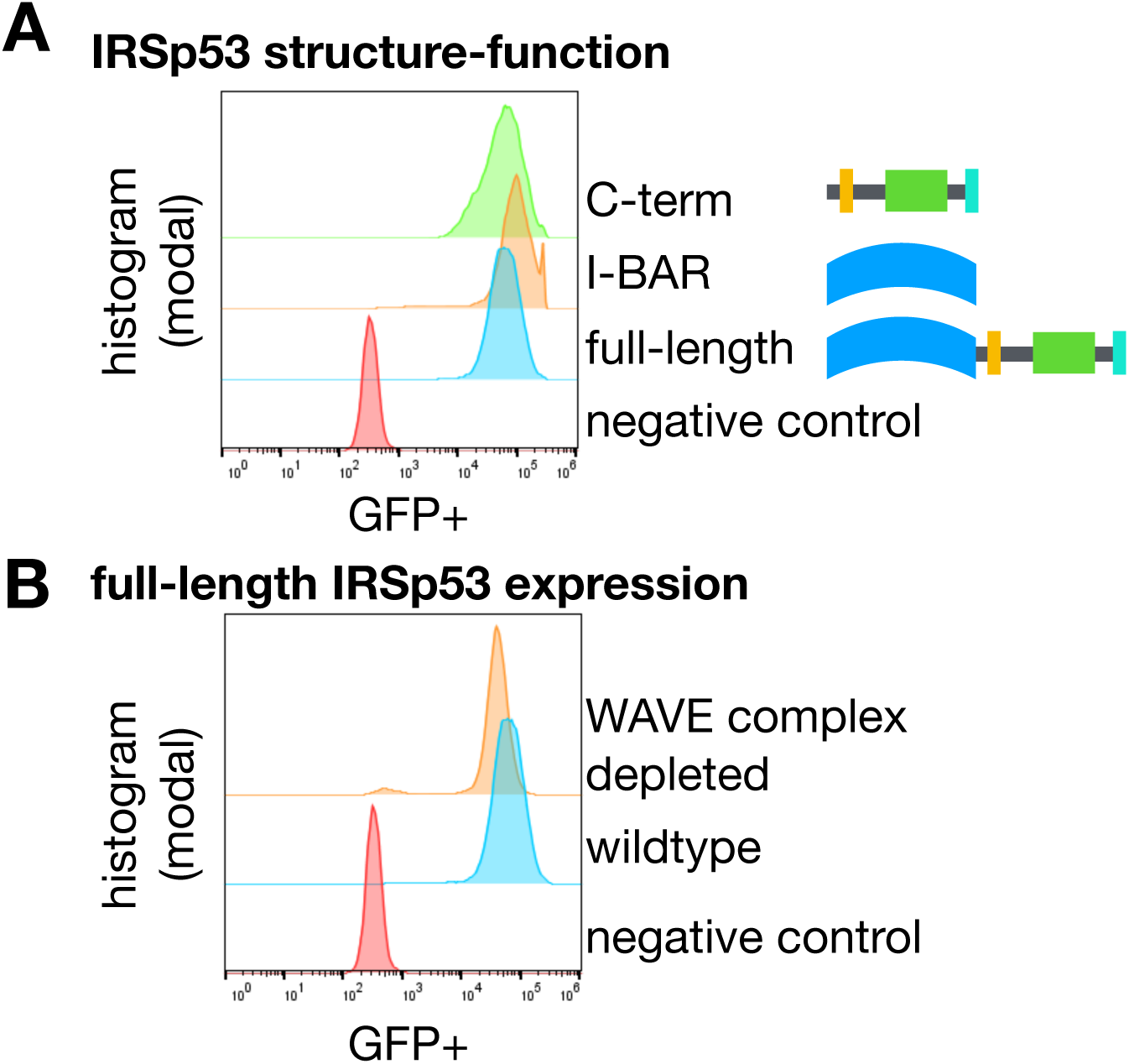
IRSp53 expression levels are comparable across conditions. **(A)** IRSp53 structure-function constructs have comparable expression levels in wildtype cells. Flow cytometry data graphed with FlowJo. **(B)** Full-length IRSp53 has comparable expression levels in wildtype and WAVE complex-depleted HL60s. Flow cytometry data graphed with FlowJo.

## SUPPLEMENTAL VIDEOS

1. **S1 Video. Lattice light sheet imaging of WAVE complex at ruffling cell edges.** dHL60 cell expressing Hem1-eGFP (green) continually exposed to chemoattractant (25nM fMLP) imaged with lattice light sheet at 1 frame every 4 seconds. Gray overlay represents cell boundary defined by cytosolic Hem1. This video corresponds to **Figure 1C** top right.
2. **S2 Video. TIRF imaging of WAVE complex at a protruding lamellipodium.** dHL60 cell expressing Hem1-eGFP (green) and stained with membrane dye (CellMask DeepRed; gray) continually exposed to chemoattractant (10nM fMLP). Both channels were imaged simultaneously with TIRF at 1 frame every 2 seconds. This video corresponds to **Figure 1C** bottom.
3. **S3 Video. TIRF-SIM imaging of WAVE complex rings in the absence of actin polymer**. dHL60 cell expressing Hem1-eGFP continually exposed to chemoattractant (10nM fMLP) and acutely treated with latrunculin A (5μM) at time = 30 seconds. TIRF-SIM imaging at 1 frame every 2 seconds. This video corresponds to **Figure 2B**.
4. **S4 Video. TIRF imaging comparison of WAVE complex structures in the absence of actin polymer.** To create a conventional resolution comparison to highlight the requirement of super-resolution microscopy to resolve diffraction-limited WAVE complex puncta as rings, the 9 images (3 phases * 3 angles) that construct TIRF-SIM images were sum-projected. This video corresponds to **Figure 2C** and **S3 Video**.
5. **S5 Video. TIRF-SIM imaging of WAVE complex rings associating with the necks of membrane invaginations.** dHL60 cell expressing Hem1-eGFP (cyan) and stained with membrane dye (DiD; magenta) continually exposed to latrunculin B (500nM). TIRF-SIM imaging at 1 frame every 10 seconds. This video corresponds to **Figure 3A**.
6. **S6 Video. ChimeraX rendering of a transendothelial cell macroaperture (TEM) tunnel saddle point.** A fixed HUVEC cell treated with ROCK inhibitor Y27632 (50μM) and labeled with a membrane dye (CellMask DeepRed). Z-stack was imaged with spinning disk confocal and rendered in ChimeraX. The video approaches a TEM and then enters the cell to face the TEM head on. This video corresponds to **Figure 4B**.
7. **S7 Video. Actin localization at TEMs.** HUVEC cell expressing actin-eGFP and treated with ROCK inhibitor Y27632 (50μM) imaged with spinning disk confocal at 1 frame every 6 seconds. This video corresponds to **S5 Figure**.
8. **S8 Video. TIRF-SIM imaging of the sufficiency of IRSp53 to form ring structures in the presence of F-actin.** dHL60 cell with high expression of IRSp53-eGFP continually exposed to chemoattractant (10nM fMLP) imaged with TIRF-SIM at 1 frame every 5 seconds. This video corresponds to **Figure 5C**.
9. **S9 Video. TIRF-SIM imaging of IRSp53 and WAVE complex colocalization at a protruding lamellipodium**. dHL60 cell expressing IRSp53-eGFP (cyan, left) and Hem1-mCherry (magenta, middle) continually exposed to chemoattractant (10nM fMLP) imaged with TIRF-SIM at 1 frame every 5 seconds. Merged channels (right) show colocalization at lamellipodium edge. This video corresponds to **Figure 5F** top.
10. **S10 Video. Full-length IRSp53 localization at TEMs.** HUVEC cell expressing full-length IRSp53-eGFP and treated with ROCK inhibitor Y27632 (50μM) imaged with spinning disk confocal at 1 frame every 2 seconds. This video corresponds to **Figure 6E** left.
11. **S11 Video. IRSp53’s I-BAR domain at a TEM.** HUVEC cell expressing I-BAR-eGFP and treated with ROCK inhibitor Y27632 (50μM) imaged with spinning disk confocal at 1 frame every 5 seconds. This video corresponds to **Figure 6E** right.
12. **S12 Video. WAVE complex localization at closing TEMs.** HUVEC cell expressing eGFP-Nap1 and treated with ROCK inhibitor Y27632 (50μM) imaged with spinning disk confocal at 1 frame every 5 seconds. This video corresponds to **Figure 7B**.

